# Standardisation of cell-free DNA measurements: An International Study on Comparability of Low Concentration DNA Measurements using cancer variants

**DOI:** 10.1101/2023.09.06.554514

**Authors:** Daniel Burke, Alison S. Devonshire, Leonardo B. Pinheiro, Gerwyn M. Jones, Kate R. Griffiths, Ana Fernandez Gonzalez, Michael Forbes-Smith, Jacob McLaughlin, Kerry R. Emslie, Christopher Weidner, Joachim Mankertz, John E. Leguizamon, Marcelo Neves de Medeiros, Roberto Becht Flatschart, Antonio M. Saraiva, Paulo Jose Iwakami Beltrao, Carla Divieto, Laura Revel, Young-Kyung Bae, Lianhua Dong, Chunyan Niu, Xia Wang, Sasithon Temisak, Sachie Shibayama, Burhanettin Yalcinkaya, Muslum Akgoz, Rainer Macdonald, Annabell Plauth, Jim Huggett

## Abstract

For the impact of genomic testing from liquid biopsies to be maximized, mechanisms to ensure reproducible and comparable test performance will be required. This can be established and maintained through reference measurement procedures and materials with property values that are internationally comparable through traceability to a common standard. To achieve this objective, an interlaboratory study was organised to explore digital PCR (dPCR) for standardisation of cell-free DNA (cfDNA) quantification.

Blinded samples of wild-type/variant mixtures of two DNA sequences (*BRAF* p.V600E single nucleotide variant or *EGFR* exon 19 deletion) were provided to 12 laboratories. Laboratories independently designed and applied dPCR assays to determine absolute and relative quantities, with no guidance provided to harmonise the approach.

The mean and coefficient of variation (CV) of copy number concentrations for variant sequences were 18 copies/μL (CV 7.2%) (*BRAF* variant sample) and 9 copies/μL (CV 25%) (*EGFR* variant sample) while the mean variant allele frequencies (vAF) were 8.0% (CV 5.3%) and 0.080% (CV 29%) respectively.

This study demonstrated that dPCR was capable of exceptional technical accuracy for variant copy number concentration and vAF, even when different assays and platforms were used. This implies that dPCR offers a unique analytical methodology that can be deployed globally in supporting comparability for cfDNA testing based on the existing framework of the International System of units of measurement.

## INTRODUCTION

The use of liquid biopsies for measuring cell-free circulating tumour DNA (ctDNA) in blood specimens has the potential to transform diagnosis of solid tumours and to monitor residual disease during treatment ^1, 2^. While the potential applications for these measurements is established, the American Society of Clinical Oncology and the College of American Pathologists concluded that there is yet insufficient evidence of clinical validity and utility for the majority of ctDNA assays in advanced cancer ^3^. Furthermore, they reported little evidence of clinical validity in early-stage cancer detection, treatment monitoring, or residual disease detection outside of clinical trials, however evidence may emerge from the clinical trials currently underway ^3^.

A pre-requisite for establishing clinical validity is analytical validity and these authors and others ^4^ recognize that studies of analytical validity need to consider routes for improved standardization to provide testing confidence. This in turn would benefit from reference systems including defined samples, reference materials with known variants at defined quantities and variant allele frequency (vAF), and the reference measurement procedures (RMPs) to characterize them. Regulatory and standards organisations have also produced guidelines and documentary standards defining the requirements for reliable clinical measurements including the use of reference materials ^5^. The lack of standardization and fact that reference systems for genetic testing are in their infancy could hinder the translation of diagnostics based on cfDNA ^6, 7^ and may be part of the reason be why the potential benefits of using cfDNA are yet to be maximized ^8^. The development of RMPs will likely assist the application of new *in vitro* diagnostics (IVD) tests using liquid biopsy samples.

Digital PCR (dPCR) has been proposed as a primary RMP that is potentially traceable to the International System of Units (SI) for quantification of KRAS proto-oncogene (*KRAS*) single nucleotide variants (SNVs) with output in concentration (copies per microliter, copies/μL) and its trueness/accuracy validated through comparison with orthogonal SI-traceable methods ^9, 10^. Whilst the performance of single dPCR assays have been validated as RMPs for *KRAS*, epidermal growth factor receptor (*EGFR*) and B-Raf proto-oncogene, serine/threonine kinase, (*BRAF*) sequence variants ^9, 11^, the degree of equivalence when using different dPCR primer/probe systems to the same sequence when applied independently by laboratories has not been evaluated. If dPCR were able to provide high interlaboratory agreement in variant quantification when alternative assays were deployed, this could have wide-ranging implications for the development of an international reference system which can be applied in multiple jurisdictions for calibration and regulation of genetic testing.

The objective of this study (‘CCQM-P184’) was to evaluate the concordance between 12 international laboratories of dPCR measurements of two actionable cancer biomarkers using study materials containing target sequences at concentrations that have been found in cfDNA extracts ^12^. Each participant developed and validated their own assays for the two cancer biomarkers; one was a SNV in *BRAF* exon 15 (1799T>A) that is a biomarker for vemurafenib therapy in malignant melanoma ^13^ and the other a 15 base pair deletion in *EGFR* exon 19 which is a selective biomarker for treatment with EGFR inhibitors ^14^. Two study materials were produced containing low concentrations (<20 copies/μL) of the *BRAF* and *EGFR* sequence variants, mimicking ctDNA concentrations in plasma extracts. Additionally, the EGFR study material was designed to have a vAF close to the value often claimed to be the limit of detection for NGS methods ^15^.

## MATERIALS AND METHODS

### Study materials

Two study materials were distributed to participants, Study Material 1 supplied by the National Measurement Institute, Australia (NMIA) (Coordinating Laboratory 1), and Study Material 2 supplied by National measurement laboratory (NML) (Coordinating Laboratory 2).

Human *BRAF* gene (GRCh37.p13, NC_000007.13 (140415749..140624564) has a SNV located in exon 15 (NM_004333.6:c.1799T>A, amino acid mutation BRAF p.V600E, Genomic Mutation ID COSV56056643). Study Material 1 consisted of a buffered solution containing a synthetic linearised plasmid in a background of sonicated human genomic DNA (gDNA) and with yeast total RNA at 40 ng/μL added as carrier.

Study Material 2 consisted of a buffered solution containing a synthetic linearised plasmid in a background of sonicated human gDNA. The plasmid included a 631 bp sequence comprising exon 19 of the human *EGFR* gene with a 15 base pair deletion (NM_005228.5:c.2236_2250del15; (Genomic Mutation ID COSV51765066) corresponding to loss of 5 amino acids in the positions 746-750.

Details of the assays used for characterization of Study Materials, for evaluation of homogeneity and storage stability, for preparation of the high concentration validation solution and for preparation of the human gDNA used for the wild-type template are given in the online Supplementary Information.

Study Materials 1 and 2 were distributed to 11 laboratories and examined blind by the two coordinating laboratories (totaling 13 participating laboratories). Participants where provided the target sequences and had to select or develop their own assays.

Measurands were defined to comply with the International Vocabulary of Metrology (VIM) ^5^ and the Guide to the Expression of Uncertainty in Measurement ^16^. Participants were requested to submit the values of three measurands of each study material:

1. The copy number concentration of the variant in copies per μL (copies/μL).
2. The copy number concentration of the reference (wild-type) sequence in copies per μL (copies/μL).
3. The ratio of the variant concentration to the sum variant and reference type concentrations (vAF).

### Assay information

Participants were advised that assay amplicon lengths should be less than 80 bp for Study Material 1 and less than 120 bp for Study Material 2. Details on the range of assays deployed by the participating laboratories are available in the online Supplementary Information Tables S11 and S12. The dPCR instrument, reagents and partition volumes used by participants are presented in Tables S13 to S14.

### Result submission and data analysis

In total, 13 participants reported results, but one was excluded for compliance reasons. One participant submitted two data sets for Study Material 1 and three participants submitted two data sets for Study Material 2. Results as submitted were curated before statistical analysis as follows: participants that did not use the sum of variant and wild-type concentrations for vAF calculation were requested to submit ratios using the sum; and each participant nominated a single set of results for each Study Material for statistical analysis (12 in total).

The participant results were compared with the coordinators’ reference values by calculating the difference between the assigned and participant average values (arithmetic mean and median). The uncertainty in this difference (*U*_Diff_) was calculated as per Equation 1:

Equation 1: Calculation of the uncertainty in the difference between coordinator’s reference value and participant average values.

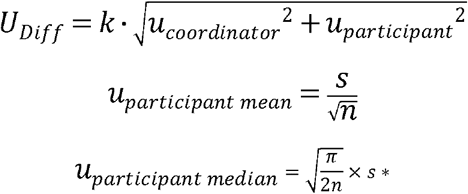

where *k* is the coverage factor corresponding to 95% confidence (*k* = 2), *u*_coordinator_ is the standard uncertainty of the coordinator’s reference value and *u*_participant_ is the standard uncertainty of the participant average value, *s* is the standard deviation of the participant average values, *n* is the number of laboratories and *s\** is the scaled median absolute deviation (MADe) which is an approximation of standard deviation for the median (calculated as the median absolute deviation (MAD) × 1.483).

## RESULTS

### Submission of results

Results from 12 laboratories for *BRAF* and *EGFR* variant and wild-type copy number concentration and vAF are shown in Figure 1 and Tables S15-S16. Eleven participants independently designed and validated dPCR assays and two worked together but submitted independently measured results. Ten participants used the QX100/200 dPCR system (Bio-Rad) and two participants QuantStudio 3D dPCR system (Thermo Fisher Scientific), with applied partition volumes ranging from 0.72-0.87 nL. Assay design factors that varied between laboratories included type of duplex assay ^17^, amplicon size and position relative to the exon that contained the variant sequence (Figure 2).

**Figure 1.**
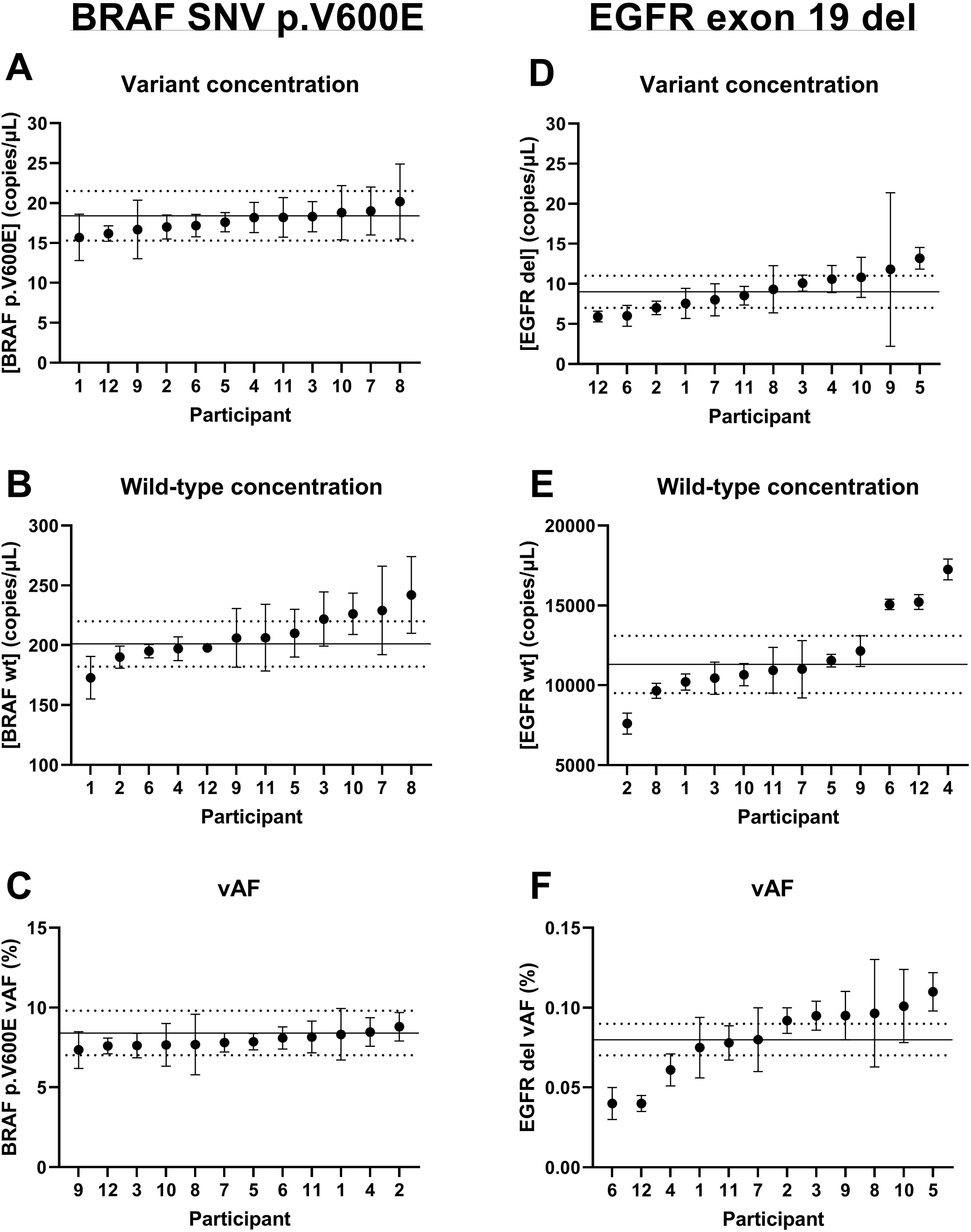
CCQM P184 study participant results. Participant results are shown in ascending order of measurand value with error bars indicating expanded uncertainty reported by participants (95% confidence). The solid and dotted lines on each graph are the coordinating laboratory’s reference value and expanded uncertainty respectively.

**Figure 2.**
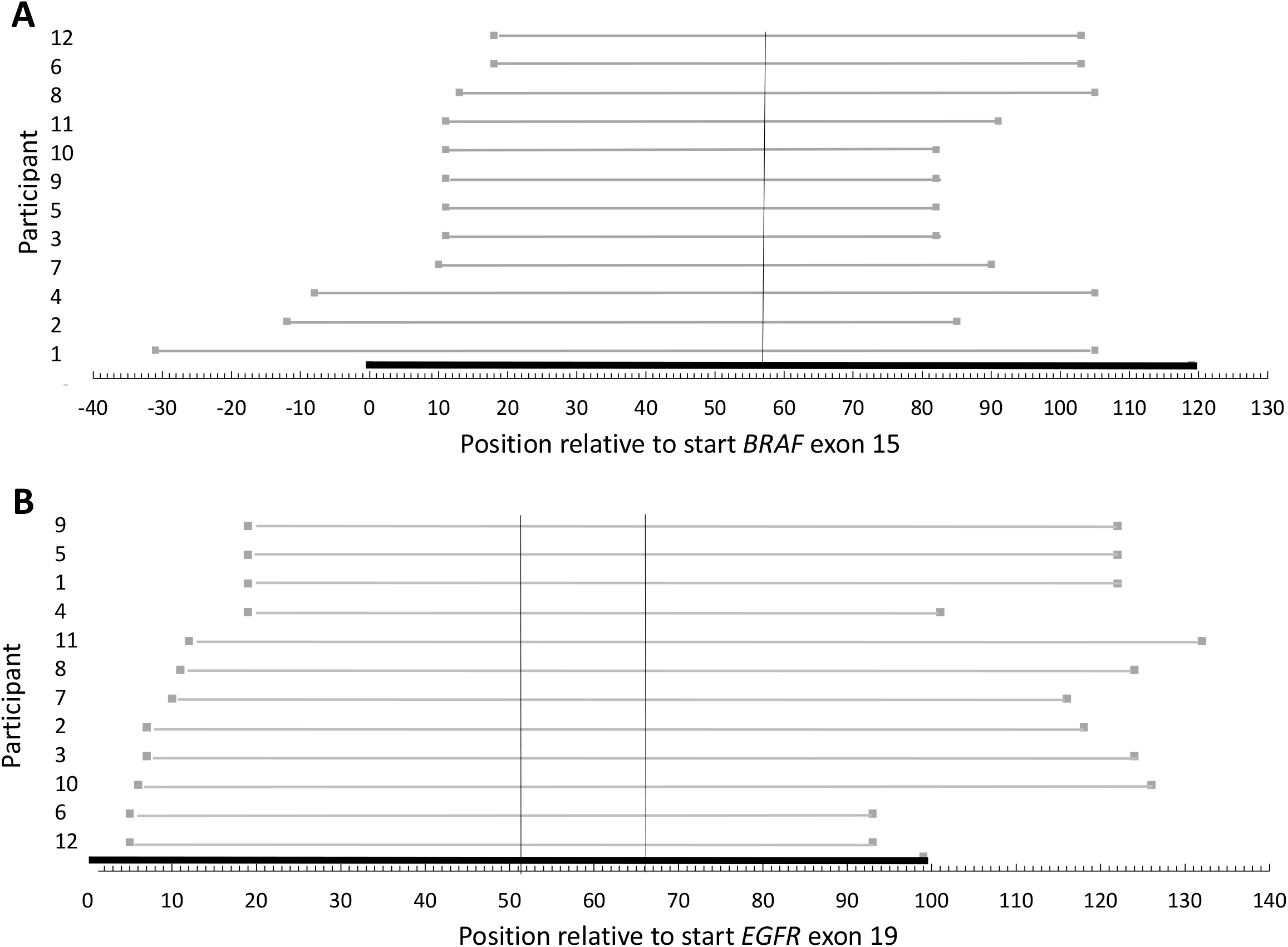
Alignment of Participant assays for Study Material template sequences. The grey lines show the length of the amplicon produced in the assay and its position relative to other participants and to A) BRAF exon 15 (the thick black horizontal line represents exon 15 and the vertical black line represents the position of the T>A mutation.) or B) EGFR exon 19 (the thick black horizontal line represents exon 19 and the vertical black lines represent the position of the 15-nucleotide deletion).

### Reproducibility

Reproducibility was evaluated by calculation of SD and non-parametric equivalent (MADe), by comparison to reference values provided by the coordinating laboratories and by visual inspection of the sorted results presented in Figure 1 as recommended in ISO 13528 ^18^. Results were compared with the coordinators’ reference values by calculating the difference and the uncertainty of the difference (*U*_Diff_) as given in Materials and Methods. Tables 1 and 2 show differences between the coordinators’ reference values and the simple and robust averages of interlaboratory results were not statistically significant.

**Table 1.** Analysis of participant results for Study Material 1.

| Measurand | Estimate | $x$ | $s$ | $u$ | $Diff$ | $U_{diff}$ |
| --- | --- | --- | --- | --- | --- | --- |
| *BRAF* p.V600E variant copy number concentration | Coordinator | 18.4 | NA | 1.50 | NA | NA |
|  | Participant mean | 17.8 | 1.3 | 0.37 | 0.6 | 3.1 |
|  | Participant median | 17.9 | 1.3 | 0.48 | 0.5 | 3.1 |
| *BRAF* wild-type copy number concentration | Coordinator | 201 | NA | 9.0 | NA |  |
|  | Participant Mean | 208 | 19 | 5.6 | 6.9 | 21.2 |
|  | Participant median | 206 | 20 | 7.2 | 5.2 | 23.1 |
| *BRAF* p.V600E vAF (%) | Coordinator | 8.40 | NA | 0.63 | NA |  |
|  | Participant mean | 7.95 | 0.43 | 0.12 | 0.45 | 1.3 |
|  | Participant median | 7.83 | 0.37 | 0.13 | 0.57 | 1.3 |

**Key** $x$ : value of measurand, variant and wild-type values are in copies/ μL, ratio values are in copies/total copies, expressed as a percentage. $s$ : standard deviation (mean value); MADe (median value) $u$ : standard uncertainty calculated as per Equation 1 $Diff$ : absolute difference from coordinator's value and participant mean or median $U_{diff}$ : expanded uncertainty of the difference between coordinator's value and participant mean or median (95% confidence, coverage factor ( $k$ ) = 2) $U_{diff} > |Diff|$ indicates that the interlaboratory study participants' average result is consistent with the coordinator's assigned value. NA: not applicable

**Table 2.** Analysis of participant results for Study Material 2.

| Measurand | Estimate | $x$ | $s$ | $u$ | $Diff$ | $U_{diff}$ |
| --- | --- | --- | --- | --- | --- | --- |
| EGFR p.Δ746-750<br>variant copy number<br>concentration | Coordinator | 8.69 | NA | 0.7 | NA | NA |
|  | Participant<br>Mean | 9.07 | 2.3 | 0.66 | -0.38 | 2.0 |
|  | Participant<br>median | 8.93 | 2.6 | 0.95 | -0.24 | 2.4 |
| EGFR wild-type copy<br>number concentration | Coordinator | 11300 | NA | 570 | NA | NA |
|  | Participant<br>Mean | 11809 | 2725 | 787 | -509 | 1948 |
|  | Participant<br>median | 11000 | 1435 | 519 | 300 | 1543 |
| EGFR p.Δ746-750<br>vAF (%) | Coordinator | 0.0772 | NA | 0.0023 | NA | NA |
|  | Participant<br>Mean | 0.0803 | 0.023 | 0.0066 | -0.0031 | 0.014 |
|  | Participant<br>median | 0.0860 | 0.016 | 0.0058 | -0.0088 | 0.013 |

**Key** $x$ : value of measurand, variant and wild-type values are in copies/μL, ratio values are in copies/total copies, expressed as a percentage. $s$ : standard deviation (mean value); MADe (median value) $u$ : standard uncertainty calculated as per Equation 1 $Diff$ : absolute difference from coordinator’s value and participant mean or median $U_{diff}$ : expanded uncertainty of the difference between coordinator’s value and participant mean or median (95% confidence, coverage factor ( $k$ ) = 2) $U_{diff} > |Diff|$ indicates that the interlaboratory study participants’ average result is consistent with the coordinator’s assigned value. NA: not applicable

Study Material 1 *BRAF* results (Measurands 1.1-1.3) for all participants based on their expanded uncertainties were within the coordinator’s uncertainty ranges (Figure 1A-C). Interlaboratory reproducibility (CV) was 7.2%, 9.3% and 5.3% for variant- and wild-type copy number concentration, and vAF results respectively (Table S15).

Study Material 2 *EGFR* deletion variant copy number concentration (Measurand 2.1) results for all participants were either within the coordinator’s reference uncertainty range, or had values close to the reference interval (Figure 1D), with a coefficient of variation of 25%. The %CV of wild-type copy number concentration (Measurand 2.2) results was 23%, and there were four results that were outside the reference range leading to the investigation described below. The vAF results for Study Material 2 (Measurand 2.3) had a CV of 29%. The mean vAF for Study Material 2 (0.08%) was about 100 times lower than for Study Material 1 (7.95%), due to a lower variant concentration combined with a high concentration of wild-type DNA (1.1 × 10^4^ copies/μL).

The sorted results (Figure 1) indicated that there may be outliers in Study Material 2 variant (participant 5) and wild-type measurements (participants 2, 4, 6 and 12), so to investigate the association with methodological factors, these results were examined further.

### Biases

Dispersion in reported variant concentration for Study Material 2 (Measurand 2.1) compared to reported measurement uncertainties using chi-squared analysis indicated that the variation between participants was not fully explained by the individually estimated uncertainties.

For Study Material 2 *EGFR* variant copy number concentration (Measurand 2.1), two laboratories (participants 5 and 9) showed a positive bias which was associated with assay format. The magnitude of the uncertainty reported by laboratory 9 (relative expanded uncertainty of 82%) was also higher than that of other participants. Instead of the competitive probe format deployed by the other participants, these laboratories opted for a “drop-off” assay with a universal reference probe and a second probe to the wild-type sequence, which can detect alternative exon 19 deletions ^19^. Both participants reported difficulty in objectively setting the threshold between positive and negative partitions in dPCR due to the proportionately high number of partitions with fluorescence intensities close to the negative population (rain) or between the double positive and single positive populations (blue and red circles, Figure S6). For these analyses, variant measurements were made more challenging due to large number of partitions in the double positive cluster (orange, Figure S6), likely to contain both wild-type and variant molecules, due to high wild-type concentration. Therefore, variant concentration and vAF could not be directly calculated based on counts in the single positive cluster (green, Figure S6).

For Study Material 2 wild-type measurement (Measurand 2.2), the three highest and the lowest results were not within the reference uncertainty interval. The three highest results were from assays with a short amplicon size (82-88 bp) and were 1.4-1.7-fold higher than the mean results for the other nine participants and consequently their vAF results were 1.5-to 2.3-fold lower. The possible bias due to amplicon size was evaluated by Coordinating Laboratory 2 using six assays of varying amplicon size and showed a clear inverse relationship between amplicon size and copy number concentration for Study Material 2, while no relationship was observed with gDNA that was not sheared (Figure 3).

**Figure 3.**
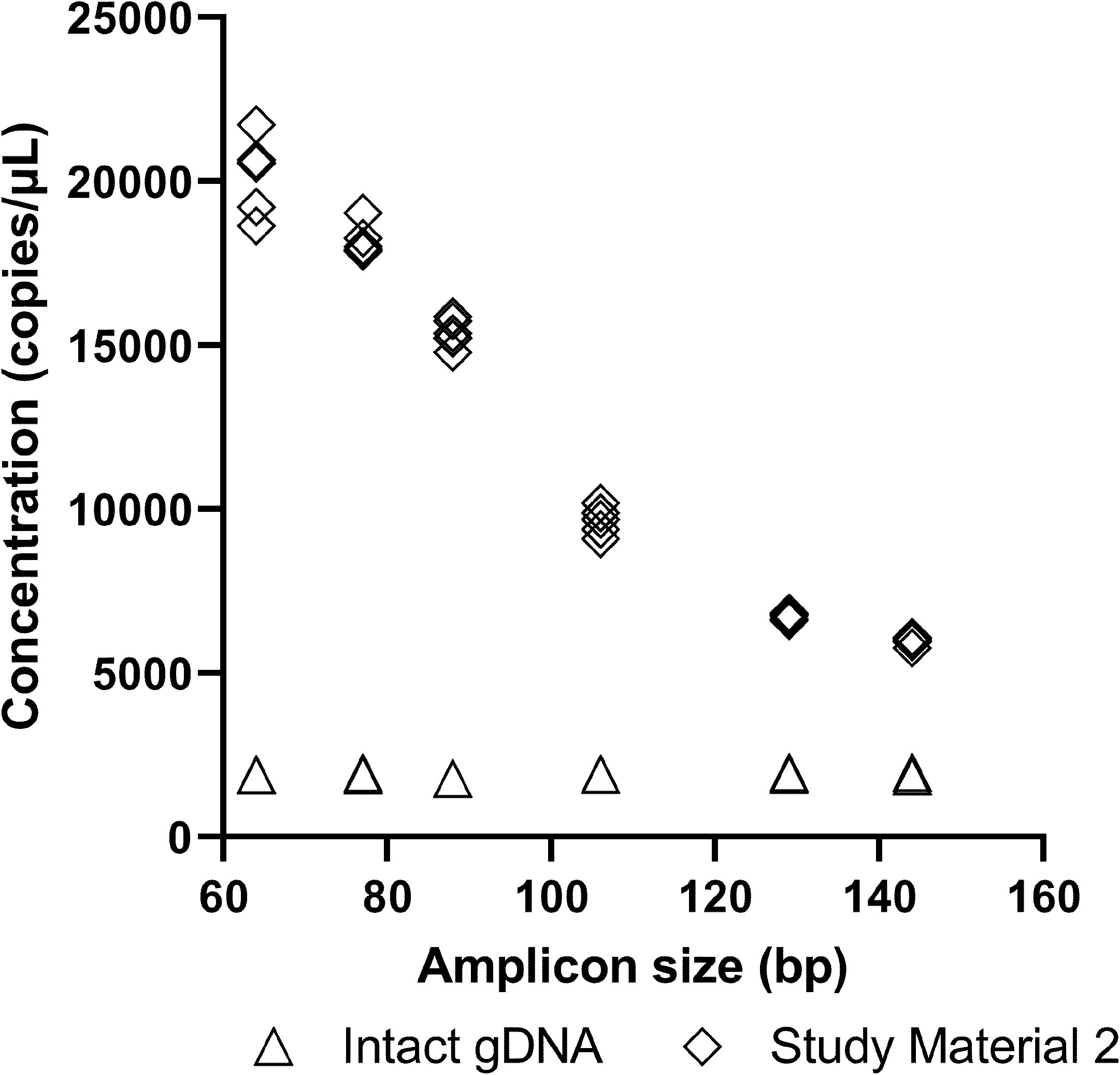
Influence of amplicon size on *EGFR* wild-type DNA copy number concentration. The impact of template fragmentation and assay amplicon size was evaluated by analysis of Study Material 2 (containing sonicated human gDNA) (*n* = 6) and intact human gDNA (*n* = 2) with six assays of varying amplicon size. Datapoints reflect individual measurements.

## DISCUSSION

This work evaluated the quantitative agreement of dPCR copy number concentration measurements of two genetic variants (and their corresponding wild-type sequences) that are used for informing treatment options in cancer. This work differs from preceding studies investigating dPCR as a reference measurement procedure since participants in this study were given the target sequence only, without a recommended measurement method. Participants had to select their own assays and were not provided with calibration materials to harmonise the approach. Therefore, this work evaluates dPCR in way that reflects current practice and includes assay selection and variation as a potential source of systematic error. It also demonstrated the participant metrology laboratories’ expertise in deploying dPCR as a molecular method per se but also for minority variant measurements.

The design of this study also provides evidence for the application of dPCR in value assignment of low vAF materials which in turn can support the establishment of reproducible IVD limits of detection and regulation of clinical tests for early cancer detection or monitoring residual disease. The narrow range of results for the variant measurements indicated that even at the low concentrations found in ctDNA, measurements may be reproducible, despite the variety of assays used. This provides evidence of the suitability of dPCR to form a part of a reference system for cancer variant measurements at low concentrations on the basis of copy number units. The SI system was initiated to improve global comparability of measurements through standard units of measurement, with enumeration of macromolecular entities such as DNA now being recognized as a dimensionless quantity in this system ^20, 21^. Our results indicate that global comparability of quantitative genetic measurements is achievable when sources of error in RMPs have been evaluated.

Two sources of bias were identified in this study that led to results being outside the consensus data set. Firstly, the “drop-off” assay format ^19^ was associated with a positive bias for *EGFR* exon 19 deletion measurements compared to the competitive format with specific probes to variant and wild-type ^17^ and suggests that the “drop-off” approach is not suitable where total DNA concentration is much higher than the variant concentration (producing average copy per partition > 2) due to the increased uncertainty in definition of variant-positive partitions.

Secondly, for *EGFR* wild-type copy number concentration (Measurand 2.2), a measurement bias was present due to different fragment lengths of the sheared gDNA used for wild-type background (Figure 4). The 1.6-fold difference in concentration measurements observed by the assays with amplicon sizes of 88 bp and 106 bp within the coordinating laboratory was consistent with the differences observed between laboratories. The wild-type template in Study Material 2 was sonicated human gDNA and it was subsequently found to have a high proportion of short fragments that may not be detected by assays with longer amplicons (Supplementary Information). For measurements where the target template corresponded to linearized plasmid or higher MW gDNA (all three *BRAF* measurands and *EGFR* variant copy number concentration), assay amplicon size and alignments showed no trends, illustrating the absence of systematic factors when measuring intact DNA. However, as a correlation between amplicon size and copy number quantities was illustrated for fragmented templates, this reflects an important consideration for both RMs using sonicated or digested genomic DNA or biological specimens where DNA fragment sizes may vary (such as for cfDNA or Formalin Fixed Paraffin Embedded).

Although this illustrates the importance of careful measurand definition for this type of study (such as the genomic coordinates of the target sequence and the source of DNA being measured), the differences are small in a biological context ^22^ and reflect factors to be considered when dealing with commutability of reference materials. This is also consistent with other studies showing that the smallest amplicons should be used for the most clinically sensitive tests ^23, 24^. While other sources of uncertainty may affect dPCR measurements such as partition volume ^25^, the magnitude of the potential variability introduced by participants applying alternative partition volumes in copy number concentration calculations was adequately covered by participants’ reported uncertainties and by the reference uncertainties provided by the study coordinators.

## Conclusion

This study has shown that independently developed dPCR assays for the quantification of genetic biomarkers gave highly concordant results through enumeration of defined DNA sequences and implies that the SI system can provide an additional route to develop global standards for genetic approaches like ctDNA testing ^3, 26^. Though dPCR may not need a calibrant, global consistency is only possible when potential sources of measurement bias have been evaluated as has occurred in this study. When dPCR measurements are accompanied by evaluation of such biases traceability to the SI is possible.

This must be undertaken during validation of candidate RMPs including testing of trueness and interlaboratory reproducibility as specified in ISO 15193. Assurance of trueness may be achieved through evaluation of systematic factors such as dPCR platform and through analysis of certified reference materials. Although the latter are limited in availability, orthogonal methods for DNA mass concentration such as isotope dilution-mass spectrometry ^27^ and gravimetrically prepared mixtures of variant and wild-type templates ^11, 22^ can support CRMs with defined DNA copy number concentration and vAF values respectively.

Additional work is required investigating these and additional sources of bias such as the method used for preparation of plasma or serum and for extraction of cell-free DNA ^28^ to improve the accuracy of such measurements. This work provides a route by which dPCR can be applied to support the application of cfDNA based diagnostics today while also offering the technological means to assist in the improvement and translate cfDNA and other molecular diagnostic solutions by providing highly accurate and reliable measurements. This outcome is also applicable to other applications where quantification of SNVs is needed such as for analysis of genome editing in food and feeds.

## Supporting information

dMIQE checklist

## Supplementary data

Additional information as noted in the text is available in Supplementary Information (containing Supplementary Tables and Supplementary Figures).

## Author Contributions

All authors confirmed they have contributed to the intellectual content of this paper and have met the following 3 requirements: (a) significant contributions to the conception and design, acquisition of data, or analysis and interpretation of data; (b) drafting or revising the article for intellectual content; and (c) final approval of the published article.

## Acknowledgements

The authors would like to thank Stephen Ellison and Simon Cowen (NML) for support with statistical analysis of study data.

