## Supplementary Information for "Standardisation of cell-free DNA measurements: An International Study on Comparability of Low Concentration DNA Measurements using cancer variants"


**AUTHORS**

Daniel Burke[1*], Alison S. Devonshire[2*], Leonardo B. Pinheiro[1], Gerwyn M. Jones[2], Kate R. Griffiths[1], Ana Fernandez Gonzalez[2], Michael Forbes-Smith[1], Jacob McLaughlin[1], Kerry R. Emslie[1], Christopher Weidner[3], Joachim Mankertz[3], John E. Leguizamon[4], Marcelo Neves de Medeiros[5], Roberto Becht Flatschart[5], Antonio M. Saraiva[5], Paulo Jose Iwakami Beltrao[5], Carla Divieto[6], Laura Revel[6], Young-Kyung Bae[7], Lianhua Dong[8], Chunyan Niu[8], Xia Wang[8], Sasithon Temisak[9], Sachie Shibayama[10], Burhanettin Yalcinkaya[11], Muslum Akgoz[11], Rainer Macdonald[12], Annabell Plauth[12] Jim Huggett[2]

**AFFILIATIONS**

1. National Measurement Institute, Australia (NMIA), Lindfield, Australia
2. National Measurement Laboratory at LGC, Teddington, United Kingdom (LGC)
3. Federal Office of Consumer Protection and Food Safety (BVL), Berlin, Germany
4. Instituto Nacional de Metrología de Colombia (INM), Bogotá, D.C, Colombia
5. National Institute of Metrology, Quality and Technology (INMETRO), Brazil
6. Istituto Nazionale di Ricerca Metrologica (INRiM), Turin, Italy
7. Korea Research Institute of Standards and Science (KRISS), Daejeon, Republic of Korea
8. National Institute of Metrology (NIM), P. R. China, Beijing, China
9. National Institute of Metrology Thailand (NIMT), Pathumthani, Thailand 12120
10. National Metrology Institute of Japan (NMIJ), Tsukuba; National Institute of Advanced Industrial Science and Technology (AIST), Ibraki, Japan
11. TUBITAK National Metrology Institute (TUBITAK UME), Gebze, Kocaeli, Turkey
12. Physikalisch-Technische Bundesanstalt (PTB), Berlin Germany

#### STUDY MATERIAL 1

##### *Preparation of Study Material 1*

A 2,136 base pair (bp) plasmid insert was designed comprising BRAF exon 15 and exon 3 together with portions of flanking intron sequences. The plasmid included a 961 bp sequence comprising BRAF exon 15 (120 bp) with a single T>A base substitution at 1799 (V600E protein mutation, COSMIC ID: COSM476), and adjacent upstream and downstream intron sequences of 392 and 449 bp, respectively. The insert was cloned into BHpUCminusMCS plasmid by Blue Heron (Bothell, WA, USA) who then supplied the the *Sca*I linearised form of the plasmid DNA (subsequently referred to as pBRAFV600E) in dried form. The linearized plasmid was dissolved in $TE_{0.1}$ and diluted with $H_2O$ to 5 ng/µL then the DNA concentration was measured by isotope dilution mass spectrometry and dPCR as previously published [1] and results are given in Figure S1.

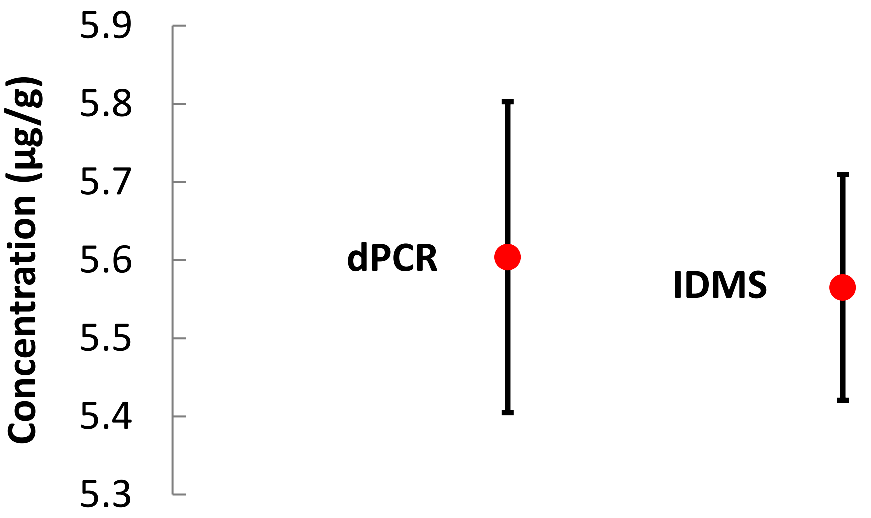


**Figure S1. Comparison of dPCR and IDMS measured BRAF V600E concentration of BRAF V600E solution used to prepare Study Material 1**. Error bars are expanded uncertainty.

Human genomic DNA (gDNA) for wild-type sequence was purified from whole blood obtained from multiple anonymous donors and stored in 10 mM Tris.HCL, 0.1 mM EDTA, pH 8.0 ($TE_{0.1}$) buffer at a concentration of approximately 228 µg/mL (Promega, WI, USA) and was fragmented by ultra-sonication using an M200 Focused-ultrasonicator (Covaris Inc, Massachusetts, USA). The sonicated material was comprised of genomic DNA fragments predominantly between 250 and 750 bp as estimated from agarose gel electrophoresis analysis shown in Figure S2.

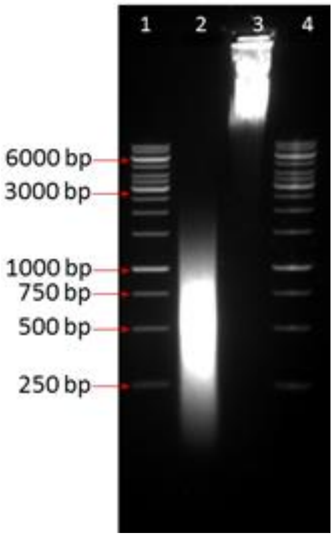


**Figure S2. Sonicated human genomic DNA used for Study Material 1.** Lane 1 & 4 O'GeneRuler 1 kb Ladder (Thermo Scientific), Lane 2 Sonicated human gDNA, Lane 3 Intact human gDNA

The sonicated human gDNA solution was diluted in sterile $TE_{0.1}$ buffer to approximately 54 ng/µL based on absorbance at 260 nm (Eppendorf Biophotometer) and mixed with sterile $TE_{0.1}$ buffer containing 40 ng/µL yeast (*S. cerevisiae*) total RNA (Sigma-Aldrich P/N R6750) to obtain a final concentration of 200 to 250 cp/µL human gDNA.

Study Material 1 was prepared by mixing 147 µL of 5,000 copies/µL pBRAFV600E with 35,000 µL of 200 to 250 copies/µL sonicated human gDNA in a background of 40 ng/µL yeast total RNA to give a nominal concentration of pBRAFV600E in Study Material 1 of approximately 21 copies/µL. Aliquots (100 µL) of Study Material 1 were dispensed into sterile 0.5 mL vials (Axygen Part SCT-050-SS-A-S) and stored at -20 ºC before transportation to participants on dry ice.

***Quantification of Study Material 1 using multiple assays***

To assess the selectivity of duplex BRAF assay 1, randomly selected vials of Study Material 1 were analysed using BRAF assay 1, pBRAFV600E construct assay and dHsaCP2000027 (Bio-Rad Laboratories Pty Ltd., Australia). Copy number concentration of plasmid pBRAFV600E was analysed using all three assays whilst copy number concentration of wild type BRAF 1799T was analysed using BRAF assay 1 and dHsaCP2000027. For each assay, three vials of Study Material 1 were analysed on separate days in replicates of five (day 1), six (day 2) or seven (day 3).

***Study Material 1 Reference value assignment and estimation of uncertainty of reference value***

For each measurand in Study Material 1, the average value for the 10 vials analysed for homogeneity assessment using duplex BRAF assay 1 was assigned as the reference value. For BRAF 1799T>A and BRAF 1799T, relative standard uncertainty for the reference value was calculated by combining relative standard uncertainties for method precision, homogeneity, short-term stability, long-term stability and droplet volume (Type B). For the ratio of BRAF 1799T>A to total BRAF 1799, relative standard uncertainty for the reference value was calculated by combining relative standard uncertainties for method precision, homogeneity, short-term stability and long-term stability. For each measurand, combined standard uncertainties were expanded to a level of confidence of 95% using a coverage factor calculated using the effective degrees of freedom obtained from the Welch-Satterthwaite equation [2].

**Table S1. Reference values for Study Material 1**

| Measurand | x | units | u | k | U | U% |
|---|---|---|---|---|---|---|
| 1.1 | 18.4 | copies/µL | 1.5 | 2.1 | 3.1 | 17% |
| 1.2 | 201 | copies/µL | 9.0 | 2.1 | 19 | 9.6% |
| 1.3 | 0.084 | Copies/total copies | 0.0063 | 2.1 | 0.014 | 16% |

x reference value, u standard uncertainty, k coverage factor for 95% level of confidence, U expanded uncertainty, U% relative expanded uncertainty

##### *Characterisation of Study Material 1*

Droplet dPCR (ddPCR) was used by the coordinating laboratory to characterise Study Material 1 and to evaluate its homogeneity and stability. Four ddPCR assays were utilised (**Table S2** and **Table S3**): (i) an inhouse designed “Competitive duplex BRAF assay 1” to BRAF variant (1799T>A) and wt (1799T) sequences, (ii) “pBRAFV600E construct assay” designed to construct specific sequences, (iii) CDKN2A Assay G, which measures a reference gene present in human gDNA, and (iv) a commercial assay “dHsaCP2000027” to BRAF variant and wt sequences.

##### *Homogeneity assessment of Study Material 1*

Evaluation of Study Material 1 homogeneity was conducted in accordance with ISO Guide 35 [3] by analysing duplicate sub-samples, each in triplicate, from 10 vials selected at random from the batch. Homogeneity testing was carried out using competitive duplex BRAF assay 1 and also using a duplex ddPCR comprising pBRAFV600E construct assay and CDKN2A Assay G (**Table S4** and **Table S5**). Copy number concentration values for each replicate were used to calculate within-vial and between-vial variances for BRAF 1799T>A and BRAF 1799T whilst ratio values for each replicate were used to calculate within-vial and between-vial variances for the ratio of BRAF 1799T>A to total BRAF 1799. The uncertainty in Study Material 1 reference values incorporates components for method precision and homogeneity taken from within-vial and between-vial variances, respectively, using competitive duplex BRAF assay 1.

**Table S2. ddPCR assays, primers and probes used by Co-ordinating Laboratory 1 for characterising plasmid BRAF V600E and Study Material 1**

| Assay | Primer/ probe | Code | Sequence (5′ - 3′) |
|---|---|---|---|
| Competitive duplex BRAF assay 1 | F | BRAF_736486 | CACCTCAGATATATTTCTTCATG |
| | R | BRAR_141236 | ATCCAGACAACTGTTCAAACTGATG |
| | P | BRAP_146853F | 6-FAM-TAGCTACAG**A**GAAATC-MGBNFQ |
| | P | BRAP_146953V | VIC-CTAGCTACAG**T**GAAATC-MGBNFQ |
| pBRAFV600E construct assay | F | BRAF_148360 | CAGGTATGCAATTTAAGGCAGGTG |
| | R | BRAR_148407 | CCATTGATGCAGAGCTAGAAACAG |
| | P | BRAP_143713H | 5-HEX-TGGAATCTCTGGGGAACGGAACTGA-BHQ-1 |
| CDKN2A Assay G | F | CDKF_855371 | GCGACTCTGGAGGACGAAG |
| | R | CDKR_852303 | TCCCCTTTTTCCGGAGAATCG |
| | P | CDKP_859975F | 6-FAM- CGCTACCTGATTCCAATTCCCCTGC-BHQ-1 |
| dHsaCP2000027 | Proprietary (Bio-Rad Laboratories Pty Ltd.) | | |

F, forward primer; R, reverse primer; P, probe; 6-FAM, 6-carboxyfluorescein; BHQ-,1 Black Hole Quencher 1; 5-HEX, hexachloro-6-carboxyfluorescein; MGBNFQ, minor groove binder non-fluorescent quencher

**Table S3: Thermal cycling conditions used by coordinating laboratory 1 for duplex assays for characterising plasmid BRAF V600E and Study Material 1**

| Assay | Cycling step | Temperature (°C) | Time | Number of cycles |
|---|---|---|---|---|
| Competitive duplex BRAF assay 1 | Enzyme activation | 95 | 10 min | 1 |
| | Denaturation | 95 | 30 sec | 45 |
| | Annealing- Extension | 59 | 60 sec (50% ramp rate) | |
| | Enzyme deactivation | 98 | 10 min | 1 |
| | Hold | 12 | ∞ | 1 |
| BRAFV600E construct assay CDKN2A Assay G | Enzyme activation | 95 | 10 min | 1 |
| | Denaturation | 95 | 30 sec | 45 |
| | Annealing- Extension | 60 | 60 sec (50% ramp rate) | |
| | Enzyme deactivation | 98 | 10 min | 1 |
| | Hold | 12 | ∞ | 1 |
| dHsaCP2000027 | Enzyme activation | 95 | 10 min | 1 |
| | Denaturation | 94 | 30 sec | 45 |
| | Annealing- Extension | 55 | 60 sec (50% ramp rate) | |
| | Enzyme deactivation | 98 | 10 min | 1 |
| | Hold | 12 | ∞ | 1 |

ANOVA of within- and between-vial variance for homogeneity data using competitive duplex BRAF assay 1 indicated no significant inhomogeneity for any measurand (*p*-values between 0.266 and 0.478). Relative standard uncertainties for homogeneity of BRAF 1799T>A, BRAF 1799T and the ratio were 3.3%, 1.2% and 0.8%, respectively (Table S). Relative standard uncertainties for method precision of BRAF 1799T>A and the ratio were 6.6% and 6.4%, respectively, due largely to stochastic effects from the relatively low copy number concentration of BRAF 1799T>A whilst relative standard uncertainty for method precision of BRAF 1799T was 2.8% (Table S). Similar results were obtained using a duplex assay comprising pBRAFV600E construct assay and CDKN2A Assay G (*p*-values between 0.613 and 0.699) (Table SS5).

**Table S4. Relative standard uncertainties for method precision and homogeneity of Study Material 1 using Competitive duplex BRAF assay 1**

| | [BRAF 1799T>A] (copies/µL) | [BRAF 1799T] (copies/µL) | Ratio[a] |
|---|---|---|---|
| Homogeneity (%) | 3.3 | 1.2 | 0.8 |
| Method precision (%) | 6.6 | 2.8 | 6.4 |

[a] BRAF 1799T>A: (BRAF 1799T>A + BRAF 1799T)

**Table S5: Relative standard uncertainties for method precision and homogeneity of Study Material 1 using a duplex assay comprising pBRAFV600E construct assay and CDKN2A Assay G**

| | [pBRAFV600E plasmid] (copies/µL) | [CDKN2A] (copies/µL) | Ratio[a] |
|---|---|---|---|
| Homogeneity (%) | 2.5 | 1.0 | 2.4 |
| Method precision (%) | 5.3 | 2.2 | 5.1 |

[a]Ratio of pBRAFV600E to sum of pBRAFV600E and CDKN2A

##### *Stability evaluation of Study Material 1*

Short-term stability of Study Material 1 was assessed using an isochronous experimental layout at both room temperature and 40 °C for 14 days, based on the initial plan to ship samples at ambient temperature. Randomly selected vials were placed at the designated temperature. At each designated time point (3, 7 and 14 days), 3 vials were taken from both room temperature and 40 °C storage and transferred to 4 ºC. On the fifteenth day, all vials which had been transferred to 4 ºC (nine vials for each incubation temperature) were analysed in triplicate (non-randomised order) by ddPCR using a duplex assay comprising pBRAFV600E construct assay and CDKN2A Assay G (**Table S2**). A *t*-test was used to determine if the slope of the linear regression over time was significantly different from zero. Standard uncertainty of the slope of the regression line (copies/µL/week) for vials stored at room temperature was used to estimate a factor for uncertainty in short-term stability over a period of one week under conditions which may be experienced during transport to participating laboratories.

Long-term stability of Study Material 1 was assessed using an isochronous experimental layout. Randomly selected vials were placed at -20 °C. At each designated time point (approximately 0, 16, 46, 76 and 166 days), 3 vials were transferred from -20 °C storage to -80 °C. After the 180 day time point, all vials which had been transferred to -80 ºC (15 vials in total) were analysed in triplicate by ddPCR using a duplex assay comprising pBRAFV600E construct assay and CDKN2A Assay G (Table S). Standard uncertainty associated with long-term stability was calculated from the standard uncertainty of the slope of the linear regression (copies/µL/month) multiplied by 12 months to incorporate the time period during which participating laboratories would complete their analysis.

No significant linear trend with time was observed for any measurand following storage of Study Material 1 at either room temperature (*p*-values between 0.191 and 0.432) or 40 °C except for the CDKN2A target which showed a significant increase with time at 40 °C (*p*-values of 0.152, 0.018 and 0.344 for plasmid BRAFV600E, CDKN2A and the ratio respectively). The observed increase in CDKN2A target over one week at 40 °C is most likely due to a small amount of evaporation from vials at elevated temperature. To ensure no significant effect during transport, vials were transported on dry ice to participating laboratories. No significant linear trend with time was observed for any measurand following long-term storage of Study Material 1 vials for 12 months at -20 °C (*p*-values between 0.320 and 0.990). Relative standard uncertainties for stability over one week at room temperature and over 12 months at -20 °C were 0.84% to 1.3% and 2.5% to 3.0%, respectively (**Table S6**).

**Table S6. Relative standard uncertainties for stability of Study Material 1**

| Condition | [BRAFV600E plasmid] (copies/µL) | [CDKN2A] (copies/µL) | Ratio[a] |
|---|---|---|---|
| One week, room temperature (%) | 1.3 | 0.84 | 1.1 |
| One week, 40 °C (%) | 3.3 | 2.9 | 3.3 |
| 12 months, -20 °C (%) | 2.5 | 3.0 | 2.7 |

Measurements were performed using BRAFV600E construct assay and CDKN2A Assay G duplex (Table S). aRatio = [BRAFV600E]/ ([BRAFV600E] + [CDKN2A])

##### *Selectivity of Assays for Quantification of Study Material 1*

To assess the selectivity of competitive duplex BRAF assay 1, randomly selected vials of Study Material 1 were analysed using competitive duplex BRAF assay 1, pBRAFV600E construct assay and dHsaCP2000027 (Bio-Rad Laboratories Pty Ltd., Australia). Copy number concentration of plasmid pBRAFV600E was analysed using all three assays whilst copy number concentration of BRAF 1799T was analysed using BRAF assay 1 and dHsaCP2000027. For each assay, three vials of Study Material 1 were analysed on separate days in replicates of five (Day 1), six (Day 2) or seven (Day 3).

Competitive duplex BRAF assay 1 targets BRAF 1799T>A and the corresponding BRAF 1799T whilst pBRAFV600E construct assay targets a synthetic junction sequence within plasmid pBRAFV600E which does not naturally occur in human gDNA. Assay dHsaCP2000027 also targets BRAF 1799T>A and the corresponding BRAF 1799T. Average copy number concentration of plasmid pBRAFV600E in ten vials randomly selected for homogeneity assessment was 18.4 copies/µL using BRAF assay 1 and in a second set of ten randomly selected vials was 18.6 copies/µL using pBRAFV600E construct assay (Table S) demonstrating excellent agreement between the two assays. Further assessment of assay selectivity was undertaken by quantifying copy number concentration of plasmid pBRAFV600E and BRAF 1799T in additional vials of Study Material 1 over three days using three and two ddPCR assays, respectively. Excellent agreement was observed between results for BRAF assay 1, dHsaCP2000027 and pBRAFV600E construct assay. Collectively, these results confirm the selectivity of BRAF assay 1 for BRAF 1799T>A.

#### STUDY MATERIAL 2

##### *Preparation of Study Material 2*

A 643 bp insert was designed to include a sequence encoding deletion mutant variant p.E746 A750del (c.2236_2250del15 COSMIC ID: COSM6335) of the human EGFR gene (GRCh38, 7:55174773..55174787; NG_007726.3: 160516 to 161161). The insert consisted of a 631 bp sequence including exon 19 of chromosome 7 with adjacent intron sequences and additional *Eco*RI and *Bam*HI restriction sites at the 5' and 3' ends, respectively. The insert was synthesised and cloned into plasmid pEX-A2 by Eurofins Genomics (Ebersberg, Germany).

The synthesised 3093 bp construct (hereon referred to as pEX-A2.EGFRΔ746-750) was linearised in-house at LGC using *Sca*I-HF restriction enzyme (NEB P/N R3122) in a reaction composed of approximately 10 ng/µL plasmid, 1x CutSmart Buffer and 0.4 Units/µL *Sca*I-HF enzyme, made up to 50 µL with Ambion nuclease-free water (Thermo Fisher Scientific P/N AM9937). The restriction digestion reaction was incubated at 37 °C for 120 minutes, followed by a 20 minutes heat inactivation at 80 °C. The digest was analysed by capillary electrophoresis (Agilent 2100 Bioanalyzer, High Sensitivity DNA kit) to confirm linearised size (Figure S3).

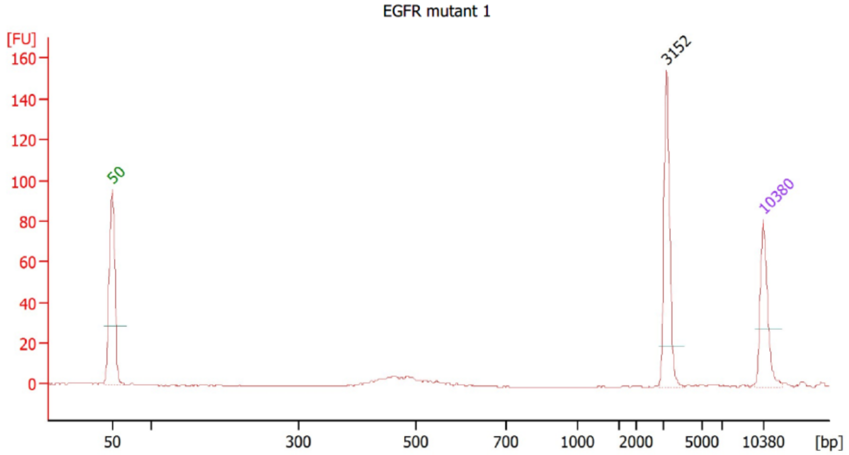


**Figure S3. Electropherogram of linearised pEX-A2.EGFRΔ746-750 (EGFR variant) analysed on the Agilent 2100 Bioanalyzer System with the High Sensitivity DNA kit.** Peaks at 50 and 10380 bp are Bioanalyzer lane size markers. Peak at 3152 bp corresponds to linearised pEX-A2.EGFRΔ746-750 (expected size 3093 bp). The sizing resolution of the High Sensitivity DNA kit is ± 20% (600-7000 bp range; Agilent Publication Number 5990-4417EN).

The linearised plasmid was then diluted gravimetrically (Mettler Toledo XP205) to ~ 1 × $10^5$ copies/µL by serial 30-fold, 10-fold and 100-fold dilution in sonicated human genomic DNA with fragment lengths between 100 and 300 base pairs as shown in Figure  (Cambio Product code: CA-972-05; ~1 × $10^4$ copies/µL based on dPCR using LGC EGFR assay (Table S7, Table S8) to produce ~ 5 mL working stock of the plasmid.

Following dPCR evaluation of the working stock, the linearised pEX-A2.EGFRΔ746-750 underwent gravimetric serial dilution to a final concentration of ~7.5 copies/µL, in sonicated human genomic DNA (Figure S4) to produce approximately 21500 µL of Study Material 2 *via* three consecutive 10-fold serial dilutions followed by a final 13.04-fold dilution, with each dilution being mixed by vortexing for 3-5 seconds. The material was subsequently divided into 50 µL aliquots in 1.5 mL sterile, nuclease-free Safe-Lock DNA Lo-bind tubes (Eppendorf) and stored at -20 °C.

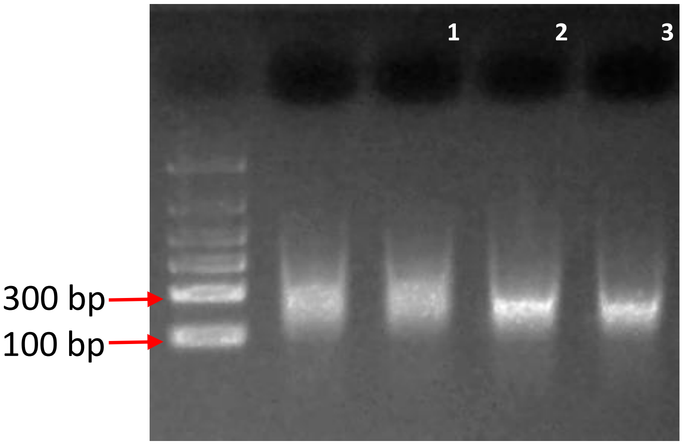


**Figure S4. Study Material 2 gDNA on 1.2% agarose gel electrophoresis (FlashGel system, Lonza).** Lane 1: FlashGel DNA QuantLadder 100 - 1500bp (Lonza; Catalogue No: 57033); Lanes 2-3: Study Material 2 (10-fold dilution); Lanes 4-5: Study Material 2 (5-fold dilution).

##### *Preparation of EGFRΔ746-750 validation solution*

A mutant only validation solution of pEX-A2.EGFRΔ746-750 was produced by volumetric serial dilution of linearised plasmid at $2.94 \times 10^9$ copies/µL to approximately $5 \times 10^5$ copies/µL in 1× TE buffer (Sigma-Aldrich P/N 93283). The resultant solution was divided into 100 × 50 µL aliquots in 1.5 mL sterile, nuclease-free Safe-Lock DNA Lo-bind tubes (Eppendorf) and stored at -20 °C.

##### *Characterisation of Study Material 2*

Characterisation of Study Material 2, including an evaluation of its homogeneity and stability was carried out by coordinating laboratory 2, using the QX200 ddPCR System (Bio-Rad) and an ‘EGFRex19del’ assay, as described in Table S7 and Table S8.

**Table S7: EGFRex19del assay used by Coordinating Laboratory 2: primer and probe sequences and concentrations.**

| Assay name | Primer/ Probe name | Sequence (5′ - 3′) | Final concentration (µM) |
|---|---|---|---|
| EGFRex19del_1 | EGFRex19del_F1 | TCCCAGAAGGTGAGAAAGTTAAAATT | 0.9 |
| | EGFRex19del_R1 | GCAAAGCAGAAACTCACATCGA | 0.9 |
| | EGFRex19del_WT1 | VIC-CAAGGAATTAAGAGAAGCA-MGB- NFQ | 0.25 |
| | EGFRex19del_MT1 | 6-FAM-CTATCAAGACATCTCCGAAA-MGB-NFQ | 0.25 |

Key: MGB, Minor Groover Binder; NFQ, Non-fluorescent quencher. Probes purchased from Thermo Fisher Scientific.

**Table S8: EGFRex19del assay used by Coordinating Laboratory 2: thermal cycling parameters.**

| Step | Time | Temp (°C) | # of cycles | Ramp rate (°C/s) |
|---|---|---|---|---|
| Enzyme activation | 10 min | 95 | 1 | 2 |
| Denaturation | 30 sec | 94 | 40 | 2 |
| Annealing/extension | 1 min | 55 | | 2 |
| Enzyme denaturation | 10 min | 98 | 1 | 2 |
| Hold (optional) | ∞ | 12 | 1 | 1 |

##### *Homogeneity assessment of Study Material 2*

For the purpose of determining a coordinator's reference value for mean concentration and evaluating the homogeneity of Study Material 2, 15 units were randomly selected from storage at -20°C and quantified by dPCR, with each analysed in replicate reactions ($n$ = 8) using the EGFRex19del assay. Samples 1-8 were analysed in a single 96-well plate and samples 9-15 in a second plate, with 8 wt only reactions included on each plate in order to calculate false positive rate for the assay. Reaction order was randomised for each plate layout.

Homogeneity results for Sample 2 were analysed using mixed effect models, allowing for variation from location (row and column), plate, and sample nested within plate. Although some marked row and plate effects were found, between-sample variability was small compared to other sources of variability, with the between-unit RSD estimated to be 0.42%, 0% and 0% for wild-type, mutant, and mutant proportion (ratio) respectively. Since between-unit variances can be small by chance, an allowance for possible inhomogeneity was calculated from the residual (within-unit) standard deviation using Equation 1.

$$u_{\mathrm{hom}} = \max\left(s_{\mathrm{b}}, \frac{s_{\mathrm{w}}}{\sqrt{n}}\right)$$

**Equation 1: Formula used to determine the uncertainty due to material inhomogeneity.** The maximum of the between-unit variance and the measurement uncertainty was applied. $u_{\mathrm{hom}}$ is the standard uncertainty associated with (possible) inhomogeneity, $s_{\mathrm{b}}$ is the estimated between-unit standard deviation, $s_{\mathrm{w}}$ the within-unit (residual) standard deviation and $n$ the number of replicates used in the homogeneity study (see ref. [4])

The resulting allowances are shown in Table S9, as relative standard deviations.

**Table S9: Study Material 2 (EGFR) homogeneity contributions (as RSD (%))**

| **Measurand** | **Variant concentration** | **Wild-type concentration** | **Variant/total copy number** |
|---|---|---|---|
| Between-unit variation (RSD) | 0.0000 | 0.0042 | 0.0000 |
| Within-unit variation (RSD) (Method precision) | 0.1904 | 0.0179 | 0.1934 |
| Homogeneity ($u_{\mathrm{hom}}$) | 0.0676 | 0.0064 | 0.0687 |
| DF | 104 | 97 | 104 |
| Homogeneity basis | Within | Within | Within |

[a]The degrees of freedom (DF) associated with $u_{hom}$are the degrees of freedom for the within- or between-unit standard deviation, depending on the basis for $u_{\mathrm{hom}}$ (see Basis).

##### *Stability Evaluation of Study Material 2*

Evaluation of the short-term stability of Study Material 2 was carried out to mimic conditions that the samples may encounter during shipping to participating laboratories. Samples were stored in dry ice, at 4 °C and 25 °C for 0 (equilibrated to storage temperature for 1 hour), 2 and 7 days, in an isochronous scheme with randomly selected replicate vials ($n$ = 6) tested at each condition/ time point combination. Samples were then stored at -20 °C overnight before being analysed by digital PCR using the EGFRex19del assay the following day. Samples were analysed across three randomized PCR plates, with samples from each temperature condition tested on a single plate. Each plate also included replicate units ($n$ = 6) each of a *wt* only control and Study Material 2 taken from -20 °C storage immediately prior to PCR setup. All samples were analysed in triplicate dPCR reactions. Statistical analysis used mixed effects models with maximum likelihood estimation. For the wild-type target, a significant effect of incubation time on dry ice was observed ($p$=0.0002); the estimated effect was +0.0153 copies/partition per day, or about 0.7% per day. No change in variant

copy number or ratio was observed in relation to time stored on dry ice or at 4 °C; however 25 °C incubation appeared to result in increases in copy number and ratio of $5.9\times10^{-5}$ copies/partition/day for variant copy number and $1.5\times10^{-5}$ per day for abundance, or about 4% and 2% per day respectively ($p$ = 0.0025 and $p$ = 0.035). Noting the direction of the effect and taking account of apparently poorer precision for one group of abundance results, neither change was considered compelling; nonetheless, participants were asked to keep materials at 4 °C or below.
A long-term stability testing scheme for Study Material 2 was designed to mimic the storage of the material at participating labs after shipping: subsets of material were stored at -20 °C and -80 °C after storage for 7 days in dry ice, as well as directly at -20 °C (Figure ). In order to monitor for a loss of stability over the course of the study, dPCR quantification of the samples was carried out on a monthly basis from 1-4 months post-storage using the EGFRex19del assay (n = 3 per unit), in parallel to the testing of the material by participants.

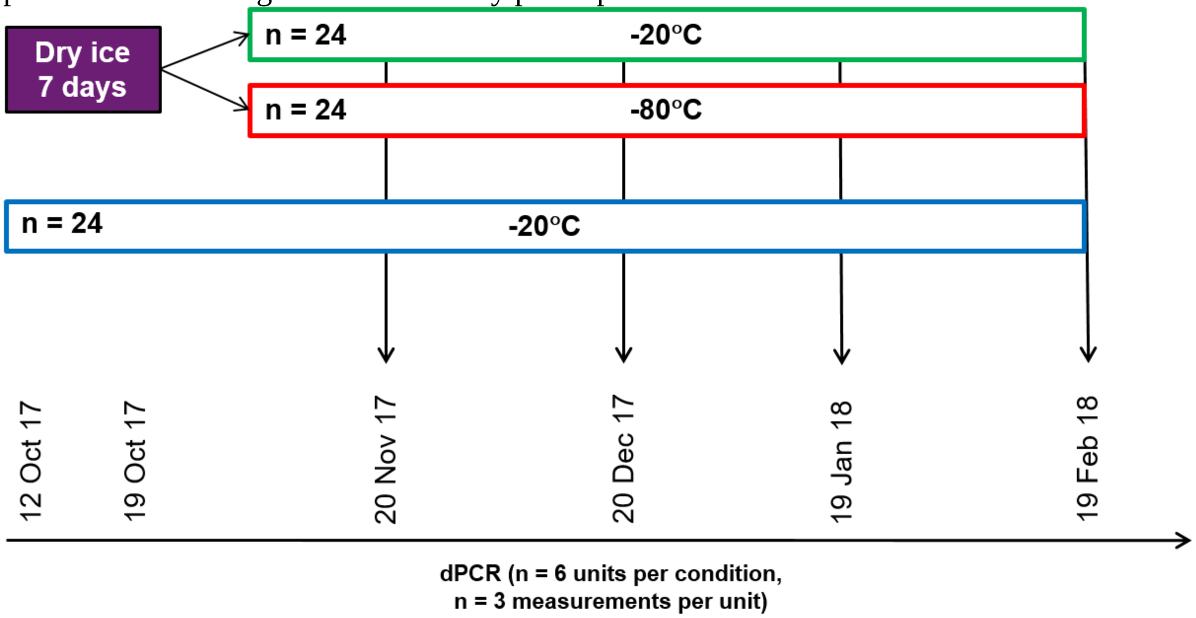


**Figure S5: Design of long-term stability study (Study Material 2).** Approximate dates when experiments were performed are indicated.

Long-term stability was assessed by analysis of lambda ($\lambda$) values. Statistical analysis used maximum likelihood treatment for mixed effects models, allowing for between-experiment and between-unit random effects. For the wild-type (after removal of two low replicates attributed to measurement issues), the effect of time was not significant but the intercept for storage at -20 °C after dry ice exposure was 0.032 copies/partition higher than for -20 °C without dry ice ($p$ = 0.005). Thus there may be a small effect associated with exposure to dry ice. For the variant target and variant abundance, no significant effects of time or temperature were found. As the effect for the wild-type target was small compared to the $\lambda$ value (c. 2 copies/partition) and there was no statistically significant effect over time for either temperature or dry ice condition, no allowance for instability was included in the measurement uncertainty of the reference values.

##### *Coordinator's reference value and uncertainty for Study Material 2*

The copy number concentration and variant copy number ratio values and uncertainty calculated from the homogeneity study of Study Material 2 were used to calculate the coordinator's reference values (Table S10). For estimating values and Type A uncertainties, models with lambda ($\lambda$) or copy number ratio as the dependent variable were fitted with the random effects: row and unit nested within plate ($\lambda_{wt}$), corresponding to measurement precision and homogeneity; or only plate ($\lambda_{mt}$ and copy number ratio), corresponding to measurement precision only. Precision associated with pipetting was captured in random effects from replicate dPCR assays; bias associated with pipetting was considered negligible. For Measurands 2.1 and 2.2, $\lambda$ values were converted to copy number concentration values applying partition volume ($V_p$) of 0.749 nL, based on the arithmetic mean of reported $V_p$ values from four laboratories (NIST, INRIM, NIB and NMIA) using ddPCR Supermix for Probes (No dUTP) (Bio-Rad) [5-7]. Partition volume uncertainty was calculated as the SD of the $V_p$ values ($u$ = 0.0385 nL; relative standard uncertainty 5.14%). Type A uncertainties were combined with the Type B uncertainty for partition volume. For Measurand 2.3, the reference value uncertainty was based on measurement precision only. The coverage factor was based on the degrees of freedom calculated using the Welch–Satterthwaite equation for combining the degrees of freedom associated with the Type A and Type B uncertainty contributions.

**Table S10. Coordinator's reference values for Study Material 2**

| Measurand | Value units | Value[a] | Standard uncertainty[b] | Coverage factor[c] | Expanded Uncertainty[a] | Relative Expanded Uncertainty[d] |
|---|---|---|---|---|---|---|
| 2.1 | copies/µL | 9 | 0.72 | 2.78 | 2 | 22% |
| 2.2 | copies/µL | $11.3 \times 10^3$ | $0.57 \times 10^3$ | 3.18 | $1.8 \times 10^3$ | 16% |
| 2.3 | copies: total copies | $8 \times 10^{-4}$ | $0.23 \times 10^{-4}$ | 2.0 | $1 \times 10^{-4}$ | 13% |

[a]Values are rounded to the same order of magnitude as expanded uncertainties. Expanded uncertainty rounded outwards to nearest whole copy (1 significant figure (s.f.)) (Measurand 2.1) and 2 s.f. (Measurand 2). Due to the very low copy number concentration of Study Material 2, the expanded uncertainty of Measurand 2.3 is also rounded to 1 s.f. (as Measurand 2.1).
[b]Standard uncertainties shown to 2 s.f. and applied unrounded to calculate expanded uncertainty.
[c]Providing level of confidence of 95%
[d]Relative expanded uncertainty shown to 2 s.f. and reflect rounded expanded uncertainties (see footnote (a)).

### STUDY PARTICIPANT PRIMER SEQUENCES AND EXPERIMENTAL DETAILS

**Table S11: Assay specifications for Study Material 1 from participants.**

| Participant | Oligonucleotide sequences and label (5′ → 3′) | Final (µM) | Amplicon size (bp) | Supplier and purification | Cycling Conditions (ramp rates for ddPCR) |
|---|---|---|---|---|---|
| 1 | Forward CTACTGTTTTCCTTTACTTACTACACCTCAGA<br>Reverse ATCCAGACAACTGTTCAAACTGATG<br>Wild type VIC-CTAGCTACAG**t**GAAATC-MGBNFQ<br>Variant 6-FAM-TAGCTACAG**a**GAAATC-MGBNFQ | 0.9<br>0.9<br>0.25<br>0.25 | 136 | ABI<br>Desalt (Pm)<br>HPLC (Pb) | 95 °C 10 min, (94 °C 30 s, 55 °C 60 s) for 40 cycles, 98 °C 10 min, hold 4 °C. ramp rate 2 °C/s |
| 2 | Forward ACTACACCTCAGATATATTTCTTC<br>Reverse TGATGGGACCCACTCCAT<br>Wild type HEX-TAGGTGATTTTGGTCTAGCTACAG**t**G-BHQ2<br>Variant FAM-TAGGTGATTTTGGTCTAGCTACAG**a**G-BHQ1 | 500 | 97 | Macrogen<br>PAGE | 95 °C 10 min, (94 °C 30 s, 57 °C 60 s) for 40 cycles, 98 °C 10 min, Cool to room temperature. Ramp rate is set to 50% of the instrument maximum speed (3.35 °C/s) ~ 1.6 °C/s. |
| 3 | Forward CATGAAGACCTCACAGTAAAAATAGGTGAT<br>Reverse TGGGACCCACTCCATCGA<br>wt VIC-CTAGCTACAG**T**GAAATC-MGBNFQ<br>variant FAM-TAGCTACAG**A**GAAATC-MGBNFQ | 0.9<br>0.9<br>0.2<br>0.2 | 71 | Thermo Fisher Scientific (primers desalted; probes HPLC-purified) | 96 °C 10 min, (98 °C 30 s, 60 °C 2 min) for 40 cycles, 60 °C 2 min, cool to 10 °C. Ramp rate: 1.5 °C/s for up rate and 1.3 °C/s for down rate. |
| 4 | Forward CACCTCAGATATATTTCTTCATG<br>Reverse ATCCAGACAACTGTTCAAACTGATG<br>wt VIC-CTAGCTACAG**T**GAAATC-MGBNFQ<br>variant 6-FAM-TAGCTACAG**A**GAAATC-MGBNFQ | 0.9<br>0.9<br>0.25<br>0.25 | 113 | primers Sigma-Aldrich HPLC<br>probes ABI HPLC | 95 °C 10 min, (95 °C 30 s,59 °C 60 s) for 45 cycles, 98 °C 10 min. Ramp rate 2.5°C/sec |
| 5 | Forward CATGAAGACCTCACAGTAAAAATAGGTGAT<br>Reverse TGGGACCCACTCCATCGA<br>wt FAM-TAGCTACAG**A**GAAATC-MGB<br>variant VIC-CTAGCTACAG**T**GAAATC-MGB<br>Reference: Oxnard et al 2014 (ref. 19) | 0.9<br>0.9<br>0.25<br>0.25 | 71 | Invitrogen<br>PAGE | 95 °C 10 min, (94 °C 30 s, 61 °C 60 s) for 40 cycles, 98 °C 10 min. Ramp rate 2 °C/s |
| 6 | Forward ACCTCACAGTAAAAATAGGTGA<br>Reverse CCAGACAACTGTTCAAACTG<br>wt VIC-AGCTACAG**t**GAAATCTC-MGBNFQ<br>variant 6-FAM-AGCTACAG**a**GAAATCTC-MGBNFQ | 0.9<br>0.9<br>0.25<br>0.25 | 85 | Metabion (Primers)<br>Thermo Fisher Scientific (Probes)<br>HPLC (Probes) | 95 °C 10 min, (95 °C 15 s, 58.4 °C 60 s) for 45 cycles, 98 °C 10 min, hold 4 °C. Ramp rate 1.5 °C/s |
| 7 | Forward CAAACTGATGGGACCCACTC<br>Reverse TCATGAAGACCTCACAGTAAA<br>wt Cal Fluor Orange 560-TCGAGATTTC**A**CTGTAGC-BHQplus<br>variant FAM-TCGAGATTTC**T**CTGTAGC-BHQplus | 0.9<br>0.9<br>0.25<br>0.25 | 80 | Biosearch Technologies<br>RP HPLC | 95 °C 10 min, (94 °C 30 s, 57 °C 60 s) for 40 cycles, 98 °C 10 min. hold 4 °C. Ramp rate 2 °C/s, except hold 1 °C/s |
| 8 | Forward TGAAGACCTCACAGTAAAAATAGGTGA<br>Reverse ATCCAGACAACTGTTCAAACTGATG<br>wt HEX-TCGAGATTTC**A**CTGTAGC-BHQplus<br>variant FAM-TCGAGATTTC**T**CTGTAGC-BHQplus | 0.4<br>0.4<br>0.3<br>0.3 | 92 | Biosearch Technologies<br>RP HPLC | 95 °C 10 min, (95 °C 30 s, 60 °C 60 s) for 40 cycles, 95 °C 10 min. Ramp rate 2 °C/s |
| 9 | Forward CATGAAGACCTCACAGTAAAAATAGGTGAT<br>Reverse TGGGACCCACTCCATCGA<br>wt HEX-CTAGCTACAG**T**GAAATC-MGBNFQ<br>variant 6-FAM-TAGCTACAG**A**GAAATC-MGBNFQ | 0.9<br>0.9<br>0.25<br>0.25 | 71 | IDT<br>Desalt (Primers)<br>HPLC (Probes) | 95 °C 10 min, (94 °C 30 s, 59 °C, 60 s) for 40 cycles, 98 °C 10 min, hold 4 °C. Ramp rates 2°C/s. |
| 10 | Forward CATGAAGACCTCACAGTAAAAATAGGTGAT<br>Reverse TGGGACCCACTCCATCGA<br>wt VIC-CTAGCTACAG**T**GAAATC-MGBNFQ<br>variant FAM-TAGCTACAG**A**GAAATC-MGBNFQ | 0.9<br>0.9<br>0.25<br>0.25 | 71 | Primer: Sentromer, Tr. (HPLC)<br>Probes: Fisher Scientific (HPLC) | 95 °C 10 min, (94 °C 30 s, 58 °C 60 s) for 39 cycles, hold 10 °C. |

| | | | | | |
|---|---|---|---|---|---|
| 11 | Forward GATCCAGACAACTGTTCAAA<br>Reverse CATGAAGACCTCACAGTAAA<br>wt HEX-ATCGAGATTTC**a**CTGTAGCT--BHQ1<br>variant FAM ATCGAGATTTC**t**CTGTAGCT--BHQ1 | 0.9<br>0.9<br>0.25<br>0.25 | 80 | Bio-Rad (HPLC) | 95 °C 10 min, (94 °C 30 sec, 55.5 °C 1 min) for 40 cycles, 98 °C 10 min, hold 4 °C. Ramp rate 2 °C/s. |
| 12 | Forward ACCTCACAGTAAAAATAGGTGA<br>Reverse CCAGACAACTGTTCAAACTG<br>wt VIC-AGCTACAG**t**GAAATCTC-MGBNFQ<br>variant 6-FAM-AGCTACAG**a**GAAATCTC-MGBNFQ | 0.9<br>0.9<br>0.25<br>0.25 | 85 | Metabion (Primers)<br>Thermo Fisher Scientific (Probes)<br>HPLC (Probes) | 95 °C 10 min, (95 °C 15 s, 58.4 °C 60 s) for 45 cycles, 98 °C 10 min, hold 4 °C. Ramp rate 2°C/s. |

**Table S12. Assay specifications for Study material 2 by all participants.**

| Participant | Oligonucleotide sequences and label (5' → 3') | Final (μM) | Amplicon size (bp) | Supplier and purification | Cycling Conditions (ramp rates for ddPCR) |
|---|---|---|---|---|---|
| 1 | Forward: GTGAGAAAGTTAAAATTCCCGTC<br>Reverse: CACACAGCAAAGCAGAAAC<br>Wild type: VIC-CTATCAAGACATCTCCGAAAG-MGB-NFQ<br>Mutant: FAM-AGGAATTAAGAGAAGCAACATC-MGB-NFQ | 0.9<br>0.9<br>0.25<br>0.25 | 103 (wt)<br>88 (mut) | Applied Biosystems. Primers (Desalt)<br>Probes (HPLC) | 95 °C for 10 min;<br>(94 °C for 30s, 55°C for 60 s) x 40 cycles;<br>98 °C for 10 min;<br>Hold 4 °C. (All ramp rates @ 2°C/s) |
| 2a | Left (F) GGATCCCAGAAGGTGAGAAA<br>Right (R) CAGCAAAGCAGAAACTCACA<br>WT probe (HEX-BHQ1)<br>ATCAAGGAATTAAGAGAAGCAACATCT<br>Mut probe (FAM-BHQ1)<br>GTCGCTATCAAGACATCTCCGAA | 0.25<br>0.25<br>0.25<br><br>0.25 | 111 (wt) | Macrogen (PAGE) | 95 °C 10:00 min<br>40 cycles of (94 °C 30 s and 57 °C 1 min), 98 °C 10 min and let cool to RT<br>Ramp rate is set to 50% of the instrument maximum speed (3.35 °C/s) ~ 1.6 °C/s. |
| 2b | As Participant 2a | 0.5 (primers)<br>0.25 (probes) | 111 (wt) | As Participant 2a | As Participant 2a |
| 3 | Forward: GGATCCCAGAAGGTGAGAAAGTT<br>Reverse: CCCACACAGCAAAGCAGAAAC<br>Wild type: VIC-AAGGAATTAAGAGAAGCAACA-MGBNFQ<br>Mutant: FAM-TCGCTATCAAGACATCT-MGBNFQ | 0.9<br>0.9<br>0.2<br>0.2 | 117 (wt)<br>102 (mut) | Thermo Fisher Scientific (primers – desalted; probes – HPLC-purified) | 96 °C for 10 min;<br>(56°C for 2min, 98 °C for 30 s) x 50 cycles;<br>56°C for 2min;<br>10°C hold.<br>Ramp rate: 1.5 °C/s for up rate and 1.3 °C/s for down rate. |
| 4 | Forward: GTGAGAAAGTTAAAATTCCCGTC<br>Reverse: ACATCGAGGATTTCCTTGTTG<br>Wild type: FAM-AGGAATTAAGAGAAGCAACATC-MGB-NFQ<br>Mutant: VIC-CTATCAAGACATCTCCGAAAG-MGB-NFQ | 0.9<br>0.9<br>0.25<br>0.25 | 82 (wt)<br>67 (mut) | Primers: Sigma-Aldrich (HPLC)<br>Probes: Thermo Fisher Scientific (HPLC) | 95 °C for 10 min;<br>(94 °C for 30 s, 56°C for 60 s) x 45 cycles;<br>98 °C for 10 min;<br>(no 4°C hold?)<br>ramp rates |
| 5 | Forward: GTGAGAAAGTTAAAATTCCCGTC<br>Reverse: CACACAGCAAAGCAGAAAC<br>Wild type: FAM-AGGAATTAAGAGAAGCAACATC-MGB<br>Reference: VIC-ATCGAGGATTTCCTTGTTG-MGB<br>Note this is a drop-off assay (Oxnard et al 2014 (ref. 19)) | 0.9<br>0.9<br>0.25<br>0.25 | 103 (wt) | Invitrogen (PAGE) | 95 °C for 10 min;<br>(94 °C for 30 s, 58°C for 60 s) x 40 cycles;<br>98 °C for 10 min<br>Ramp rate 2 °C/s |
| 6 | Forward: CTGGATCCCAGAAGGTGAGA<br>Reverse: GATTTCCTTGTTGGCTTTCG<br>Wild type: VIC-CTATCAAGGAATTAAGAG-MGBNFQ<br>Mutant: 6-FAM-CGCTATCAAGACATC-MGBNFQ | 0.9<br>0.9<br>0.25<br>0.25 | 88 (wt) / 73 (mut) | Primers: Metabion (HPLC)<br>Probes: Thermo Fisher Scientific (HPLC) | 95 °C for 10 min;<br>(95 °C for 15 s, 52°C for 60 s) x 75 cycles;<br>98 °C for 10 min;<br>4 °C hold |

| | | | | | |
|---|---|---|---|---|---|
| | | | | | ramp rate 1.5 °C/s |
| 7 | Forward: TCCCAGAAGGTGAGAAAGTTAAAATT<br>Reverse: GCAAAGCAGAAACTCACATCGA<br>Wild type: VIC-CAAGGAATTAAGAGAAGCA-MGBNFQ<br>Mutant: FAM-CTATCAAGACATCTCCGAAA-MGBNFQ | 0.9<br>0.9<br>0.25<br>0.25 | 106 (wt)<br>91 (mut) | Primers: Eurofins (HPLC)<br>Probes: Thermo Fisher Scientific (HPLC) | 95 °C for 10 min;<br>(94 °C for 30 s, 55°C for 60 s) x 40 cycles;<br>98 °C for 10 min;<br>4 °C hold<br>All ramp rates are 2°C/s, with the exception of the drop down to 4°C, which was @ 1°C/s |
| 8 | Forward: CCCAGAAGGTGAGAAAGTTAAAATTC<br>Reverse: CCCACACAGCAAAGCAGAAA<br>Wild type: FAM-AGATGTTGCTT CTCTTAATTCCTT-BHQ-1plus<br>Mutant: HEX-CTTTCGGAGATGTCT TGATAGCG-BHQ-1plus | 0.5<br>0.5<br>0.3<br>0.3 | 113 (wt)<br>98 (mut) | Biosearch Technologies<br>Primers: RPC<br>Probes: RP-HPLC | 95 °C for 10 min;<br>(95 °C for 30 s, 56°C for 60 s) x 40 cycles;<br>98 °C for 10 min;<br>4 °C hold<br>All ramp rates @ 1°C/s |
| 9 | Forward: GTGAGAAAGTTAAAATTCCCGTC<br>Reverse: CACACAGCAAAGCAGAAAC<br>Reference: HEX-ATCGAGGATTTCCTTGTTG-MGB-NFQ<br>Wild type: FAM-AGGAATTAAGAGAAGCAACATC-MGB-NFQ<br>(Note: drop-off assay (Oxnard et al 2014 (ref. 19)) | 0.9<br>0.9<br>0.2<br>0.1 | 103 (wt) | IDT (primers: desalt)<br>IDT (probes: HPLC) | 95 °C for 10 min;<br>(94 °C for 30 s, 55°C for 60 s) x 45 cycles;<br>98 °C for 10 min;<br>4 °C hold<br>Ramp rates 2°C/s |
| 10 | Forward: TGGATCCCAGAAGGTGAGAAAGTT<br>Reverse: GCAGAAACTCACATCGAGGA<br>Wild type: VIC-CCCGTCGCTATCAAGACATCTCCGA-BHQ1<br>Mutant: FAM-AAGAGAAGCAACATCTCCGAAAGCCA-BHQ1 | 0.9<br>0.9<br>0.5<br>0.25 | 120 (wt)<br>105 (mut) | Primers: Sentromer, Tr (HPLC)<br>Probes: Macrogen, Korea (HPLC) | 95 °C for 10 min;<br>(94 °C for 30 s, 58°C for 60 s) x 39 cycles;<br>10°C hold |
| 11 | Forward: CCAGAAGGTGAGAAAGTTAAA<br>Reverse: AAACTCACATCGAGGATTTC<br>Wild type: HEX-AATTAAGAGAAGCAACATCTCCG-BHQ1<br>Mutant: FAM-TCGCTATCAAGACATCTCC-BHQ1 | 0.9<br>0.9<br>0.25<br>0.25 | 120 (wt)<br>105 (mut) | BioRad (HPLC) | 95 °C for 10 min;<br>(94 °C for 30 s, 55.5°C for 60 s) x 40 cycles;<br>98 °C for 10 min;<br>4 °C hold<br>Ramp rate 2 °C/s (except Hold) |
| 12 | Forward: CTGGATCCCAGAAGGTGAGA<br>Reverse: GATTTCCTTGTTGGCTTTCG<br>Wild type: VIC-CTATCAAGGAATTAAGAG-MGBNFQ<br>Mutant: FAM-CGCTATCAAGACATC-MGBNFQ | 0.9<br>0.9<br>0.25<br>0.25 | 88 (wt) / 73 (mut) | Primers: Metabion (HPLC)<br>Probes: Thermo Fisher Scientific (HPLC) | 95 °C for 10 min;<br>(95 °C for 15 s, 52°C for 60 s) x 75 cycles;<br>98 °C for 10 min;<br>4 °C hold<br>Ramp rate 2°C/s |

**Table S13. Instrument and reagent information for analysis of Study Material 1**

| ID | Instrument | Mastermix | Thermal Cycler | Prepared (µL) | Loaded (µL) | Mean $N_T$ | Min $N_T$ | Max $N_T$ | $V_P$ (nL) | Standard uncertainty ($V_P$) (nL) | Software |
|---|---|---|---|---|---|---|---|---|---|---|---|
| 1 | QX200 | 1 | T100 | 25 | 20 | 17185 | 15359 | 19160 | 0.85 | 0.0105 | QuantaSoft v1.7.4.0917 |
| 2 | QX200 | 1 | Veriti 96-Well | 20.5 | 20 | 16863.37 | 12730 | 19943 | 0.85 | Not considered | QuantaSoft v1.7.4 |
| 3 | QS3D | 3 | GeneAmp PCR System 9700 | 60 | 15 | 17184 | 14273 | 18341 | 0.7532 | 0.0036 | QuantStudio 3D AnalysisSuite |
| 4 | QX100 | 2 | C1000, deep wells | 25 | 25 | 17478 | 11498 | 20095 | 0.792 | 0.0162 | QuantaSoft v1.7.4.0917 |
| 5 | QX200 | 1 | Veriti | 22 | 20 | 14114 | 10657 | 22452 | 0.838 | 0.0067 (0.8%) | QuantaSoft v1.7.4 |
| 6 | QX200 | 1 | Mastercycler Nexus | 220 | 20.0 | 18413 | 17659 | 19031 | 0.87 | 0.087 | QuantaSoft Analysis Pro |
| 7 | QX200 | 2 | C1000 | 22 | 20 | 16127 | 12488 | 19153 | 0.749 | 0.038 | QuantaSoft v1.6.6.0320 |
| 8 | QX200 | 1 | CFX96 Deep Well | 25 | 22 | 17296 | 15052 | 20107 | 0.819 | 0.0086 | QuantaSoft 1.7.4.0917 |
| 9 | QX200 | 2 | C1000 | 23 | 22.4 to 23 | 15680 | 11649 | 19411 | 0.786 | 0.036 | QuantaSoft v1.6.6.0320 |
| 10 | QS3D | 3 | QuantStudio™ 3D Digital PCR Instrument | 14.5 | 14.5 | 17068 | 16046 | 18486 | 0.755 | 0.023 | QuantStudio 3D AnalysisSuite |
| 11 | QX200 | 2 | TX100 | 22 | 20 | 13419 | 10515 | 19134 | 0.720 | 0.0065 | QuantaSoft, v1.7.4.0917 |
| 12 | QX200 | 1 | C1000 | 22 | 20.0 | 17638 | 14992 | 19355 | 0.87 | 0.087 | QuantaSoft Analysis Pro |

**Key:** Prepared (µL) is volume of reaction mix prepared for each replicate, Loaded (µL) is volume of reaction mix leaded for each replicate. $N_T$, total accepted partitions; $V_P$, partition volume

**Mastermix**: 1. Supermix for Probes, ddPCR Supermix for Probes (Bio-Rad Laboratories Pty Ltd.); 2. No dUTP, ddPCR Supermix for Probes (No dUTP) (Bio-Rad Laboratories Pty Ltd.); 3. QuantStudio 3D digital PCR Master Mix v2 (ThermoFisher Scientific)

**Instrument manufacturers**: AutoDG, C1000, CFX96, QX100 and QX200 (Bio-Rad); QS 3D: QuantStudio 3D Digital PCR System (Thermo Fisher Scientific); Mastercycler Nexus (Eppendorf); Veriti 96-Well and GeneAmp PCR System 9700 (Applied Biosystems)

**Table S14 Instrument and reagent information for analysis of Study Material 2**

| ID | Instrument | Mastermix | Thermal Cycler | Prepared (µL) | Loaded (µL) | Mean $N_T$ | Min $N_T$ | Max $N_T$ | $V_P$ (nL) | Standard uncertainty ($V_P$) | Analysis Software |
|---|---|---|---|---|---|---|---|---|---|---|---|
| 1 | QX200 | 1 | T100 | 25 | 20 | 16991 | 11609 | 20989 | 0.85 | 0.0105 | QuantaSoft v 1.7.4.0917 |
| 2 | QX200 | 1 | ABI Veriti 96 | 20.5 | 20.0 | 16877.7 | 10938 | 20240 | 0.85 | 1.8% | QuantaSoft 1.7.4 |
| 3 | QS3D | 3 | GeneAmp PCR System 9700 | 60 | 15 | 16763 | 16066 | 18152 | 0.7532 | 0.0036 | QuantStudio 3D Analysis Suite |
| 4 | QX100 | 2 | C1000, deep wells | 25 | 25 | 15065 | 10,004 | 19,030 | 0.795 | 1.2% | QuantaSoft v 1.7.4.0917 |
| 5a | QX200 | 1 | ABI Veriti 96 | 22 | 20 | 13241 | 11130 | 15787 | 0.838 | 0.0067 (0.8%) | QuantaSoft 1.7.4 |
| 5b | QX200 | 1 | ABI Veriti 96 | 22 | 20 | 12985 | 10722 | 16423 | 0.838 | 0.0067 (0.8%) | QuantaSoft 1.7.4 |
| 6 | QX200 | 1 | Mastercycler Nexus | 22 | 20.0 | 15239 | 12 925 | 16 451 | 0.87 | 0.087 | QuantaSoft Analysis Pro |
| 7 | QX200 | 2 | C1000 (Bio-Rad) | 22 | 20 | 15741 | 13992 | 17040 | 0.749 | 0.038 | QuantaSoft 1.6.6.0320 |
| 8 | QX200 | 1 | CFX96 Deep Well | 25 | 22 |  | 14410 | 20977 | 0.819 | 0.0086 | QuantaSoft 1.7.4.0917 |
| 9 | QX200 | 1 | C1000 (BioRad) | 24 | 22.4 to 23 uL | 16625 | 14187 | 18847 | 0.760 | 0.009 | QuantaSoft v 1.6.6.0320 and QuantaSoft Analysis Pro v 1.0 |
| 10a | QS3D | 3 | QS3D | 14.5 | 14.5 | 16274 | 15139 | 17863 | 0.755 | 0.023 | QuantStudio 3D AnalysisSuite |
| 10b | QS3D | 3 | QS3D | 14.5 | 14.5 | 16623 | 15474 | 17586 | 0.755 | 0.023 | QuantStudio 3D AnalysisSuite |
| 11 | QX200 | 2 | Tx100 | 22 | 20 | 13916 | 10515 | 19134 | 0.720 | 0.0065 | QuantaSoft V 1.7.4.0917 |
| 12 | QX200 | 1 | C1000 (BioRad) | 22 | 20.0 | 13 520 | 11 900 | 14 777 | 0.87 | 0.087 | QuantaSoft Analysis Pro |

Participants 1, 4 and 9 used Auto Droplet Generator

**Key**: see Table S15.

#### PARTICIPANT STUDY RESULTS

**Table S15: Participant Results (Study Material 1)**

| | | *Measurand 1.1 (BRAF p.V600E copy number concentration (µL$^{-1}$))* | | | | | *Measurand 1.2 (BRAF wild-type copy number concentration (µL$^{-1}$))* | | | | | *Measurand 1.3 (BRAF p.V600E vAF)** | | | | |
|---|---|---|---|---|---|---|---|---|---|---|---|---|---|---|---|---|
| **Lab ID** | ***Nominated ***** | ***x*** | ***u*** | ***k*** | ***U*** | ***Rel U (%)*** | ***x*** | ***u*** | ***k*** | ***U*** | ***Rel U (%)*** | ***x*** | ***u*** | ***k*** | ***U*** | ***Rel U (%)*** |
| **1** | | 15.69 | 1.45 | 2 | 2.91 | 18.53 | 172.83 | 8.87 | 2 | 17.73 | 10.26 | 0.0832 | 0.0081 | 2 | 0.0162 | 19.44 |
| **2** | | 17 | 0.74 | 2.00 | 1.50 | 8.60 | 190.00 | 4.60 | 2.00 | 9.30 | 4.90 | 0.0880 | 0.0042 | 2.1 | 0.0089 | 10 |
| **3** | | 18.3 | 0.9 | 2 | 1.9 | 10.3 | 221.9 | 11.3 | 2 | 22.6 | 10.2 | 0.0762 | 0.0039 | 2 | 0.0077 | 10.1 |
| **4** | | 18.2 | 0.95 | 2.011 | 1.9 | 10.60 | 197 | 5.1 | 2.001 | 10 | 5.20 | 0.0847 | 0.0044 | 2.026 | 0.0090 | 10.6 |
| **5a** | Yes | 17.6 | 0.6 | 2 | 1.2 | 7.10 | 210 | 10 | 2 | 20 | 10 | 0.0786 | 0.0026 | 2 | 0.0052 | 6.6 |
| **5b** | No | 17.7 | 0.6 | 2 | 1.2 | 6.9 | 195 | 5 | 2 | 10 | 5 | 8.16 | 0.29 | 2 | 0.58 | 7.1 |
| **6** | | 17.2 | 0.6 | 2.26 | 1.4 | 8.2 | 195 | 2.5 | 2.26 | 5.6 | 2.9 | 0.081 | 0.003 | 2.26 | 0.007 | 8.6 |
| **7** | | 19 | 1.22 | 2.4 | 3 | 16 | 229 | 11.9 | 3.11 | 37 | 16 | 0.078 | 0.00235 | 2.45 | 0.006 | 7.7 |
| **8** | | 20.2 | 2.4 | 2 | 4.7 | 23 | 242 | 16 | 2 | 32 | 13 | 0.0768 | 0.0096 | 2 | 0.019 | 25 |
| **9** | | 16.683 | 1.837 | 2 | 3.673 | 22.018 | 206.151 | 12.324 | 2 | 24.649 | 11.957 | 0.07339 | 0.005801 | 2 | 0.011602 | 15.808 |
| **10** | | 18.8 | 1.7 | 2 | 3.4 | 18 | 226.3 | 8.6 | 2 | 17.3 | 7.6 | 0.0766 | 0.0067 | 2 | 0.0134 | 17 |
| **11** | | 18.22 | 1.24 | 2 | 2.48 | 13.6 | 206.33 | 13.95 | 2 | 27.9 | 13.5 | 0.0816 | 0.0050 | 2 | 0.01 | 12.5 |
| **12** | | 16.19 | 0.46 | 2.11 | 0.98 | 6.05 | 197.74 | 1.36 | 2.11 | 2.86 | 1.45 | 0.076 | 0.002 | 2.11 | 0.005 | 6.03 |
| **Summary data***** | | **MEAN** | **SD** | **CV%** | | | **MEAN** | **SD** | **CV%** | | | **MEAN** | **SD** | **CV%** | | |
| | | 17.8 | 1.3 | 7.2% | | | 208 | 19 | 9.3 | | | 0.080 | 0.0043 | 5.3% | | |

***Key****: x, reported value; u, standard measurement uncertainty, k, coverage factor; U, expanded measurement uncertainty (95% confidence); Rel U, relative expanded uncertainty (=U/x)*
**vAF values and uncertainties are shown as numbers (not percentage)*
***Nominated result for laboratories submitting >1 result*
**** Nominated results only*

**Table S16: Participant Results (Study Material 2)**

| | | Measurand 2.1 (EGFR p.D746-750 copy number concentration ($\mu L^{-1}$)) | | | | | Measurand 2.2 (EGFR wild-type copy number concentration ($\mu L^{-1}$)) | | | | | Measurand 2.3 (EGFR p.D746-750 vAF)* | | | | |
|---|---|---|---|---|---|---|---|---|---|---|---|---|---|---|---|---|
| **Lab ID** | Nominated* | ***x*** | ***u*** | ***k*** | ***U*** | ***Rel U (%)*** | ***x*** | ***u*** | ***k*** | ***U*** | ***Rel U (%)*** | ***x*** | ***u*** | ***k*** | ***U*** | ***Rel U (%)*** |
| **1** | | 7.57 | 0.94 | 2 | 1.88 | 24.78 | 10203.06 | 254.48 | 2 | 508.97 | 4.99 | 0.000750 | 0.000090 | 2 | 0.00019 | 24.915 |
| **2a** | Yes | 7.00 | 0.40 | 2.10 | 0.84 | 12 | 7600.00 | 310.00 | 2.10 | 660 | 8.6 | 0.000920 | 0.000038 | 2.1 | 0.000080 | 8.7 |
| **2b** | No | 7.90 | 0.44 | 2.20 | 0.96 | 12 | 7500.00 | 360.00 | 2.20 | 780 | 10 | 0.001100 | 0.000046 | 2.2 | 0.000100 | 9.5 |
| **3** | | 10.10 | 0.5 | 2 | 1 | 10 | 10438 | 503 | 2 | 1005 | 9.6 | 0.000950 | 0.000045 | 2 | 0.000091 | 9.6 |
| **4** | | 10.60 | 0.86 | 2 | 1.7 | 16.30 | 17250 | 327 | 2.0 | 650 | 3.8 | 0.00061 | 0.000050 | 2.002 | 0.00010 | 16.3 |
| **5a** | Yes | 13.19 | 0.68 | 2 | 1.36 | 10 | 11540 | 197 | 2 | 393 | 3.4 | 0.0011 | 0.0001 | 2 | 0.0001 | 10 |
| **5b** | No | 22.49 | 2.86 | 2 | 5.71 | 25 | 12425 | 180 | 2 | 360 | 2.9 | 0.0018 | 0.0002 | 2 | 0.0004 | 23 |
| **6** | | 6.00 | 0.54 | 2.36 | 1.3 | 21.25 | 15074 | 137 | 2.36 | 325 | 2.2 | 0.0004 | 0.00004 | 2.36 | 0.0001 | 22.0 |
| **7** | | 8.00 | 0.655 | 2.38 | 2 | 22 | 11000 | 608 | 2.80 | 1800 | 16 | 0.00080 | 0.00004 | 3 | 0.00020 | 25 |
| **8** | | 9.32 | 1.47 | 2 | 2.94 | 31 | 9650 | 236 | 2 | 472 | 5 | 0.00097 | 0.00017 | 2 | 0.00034 | 35 |
| **9** | | 11.80 | 4.8 | 2 | 9.6 | 81.58 | 12138.7 | 486 | 2 | 972 | 8.01 | 0.00095 | 0.00008 | 2 | 0.00015 | 15.89 |
| **10a** | No | 17.7 | 1.8 | 2 | 3.7 | 21 | 12292 | 422 | 2 | 845 | 7 | 0.00138 | 0.00013 | 2 | 0.00026 | 19 |
| **10b** | Yes | 10.8 | 1.3 | 2 | 2.5 | 24 | 10661 | 347 | 2 | 693 | 7 | 0.00101 | 0.00011 | 2 | 0.00023 | 22 |
| **11** | | 8.53 | 0.58 | 2 | 1.16 | 13.6 | 10934 | 715 | 2 | 1431 | 13.5 | 0.000779 | 0.000054 | 2.00 | 0.000108 | 13.5 |
| **12** | | 5.92 | 0.312 | 2.13 | 0.67 | 11.25 | 15221.87 | 218.35 | 2.13 | 465.40 | 3.06 | 0.0004 | 0.00002 | 2.13 | 0.00005 | 12.66 |
| **Summary data*** ** | | **MEAN** | **SD** | **CV%** | | | **MEAN** | **SD** | **CV%** | | | **MEAN** | **SD** | **CV%** | | |
| | | 9.1 | 2.3 | 25% | | | 11809 | 2725 | 23% | | | 0.00080 | 0.00023 | 29% | | |

***Key**: x, reported value; u, standard measurement uncertainty, k, coverage factor; U, expanded measurement uncertainty (95% confidence); Rel U, relative expanded uncertainty (=U/x)*
**vAF values and uncertainties are shown as numbers (not percentage)*
***Nominated result for laboratories submitting >1 result*
**** Nominated results only*

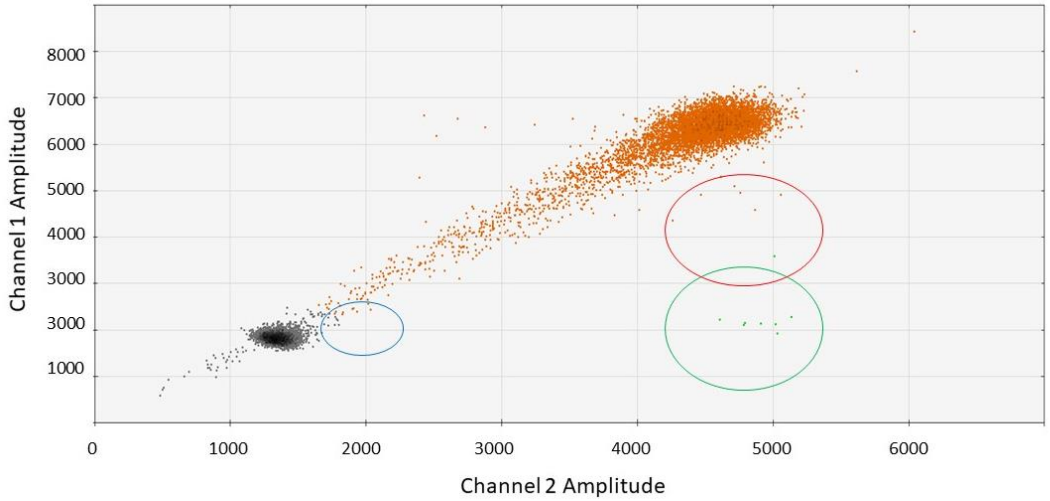


**Figure S6. dPCR 2D signal intensity plot of Study Material 2 analysed using a non-competitive assay**. The double positive population (signal from both reference and wild-type sequences is coloured orange, the single positive population (signal from only reference sequence probe) is coloured green and the double negative population is coloured black. The blue circle on the left is the intermediate fluorescence region that could be included with the single positive population if using quadrant thresholding and the red circle on the right illustrates the region that may include dual occupancy partitions, the green circle is the single positive region (containing only molecules with the variant sequence). The study material was analysed using the method specified in main text ref 19.
