## Supplementary material for "Standardisation of cell-free DNA measurements: An International Study on Comparability of Low Concentration DNA Measurements using cancer variants": dMIQE checklist

*Burke, Devonshire et al*

**dMIQE checklist**

**Key:**

Study Material 1 (SM1)

Study Material 2 (SM2)

N/A: Not applicable

| ITEM TO CHECK | PROVIDED | COMMENT |
|---|---|---|
| | Y/N | |
| **1. SPECIMEN** | | |
| Detailed description of specimen type and numbers | Y | Supplementary Information page 2 (Study Material 1 (SM1)); page 7-8 (Study Material 2 (SM2)) |
| Sampling procedure (including time to storage) | N | N/A |
| Sample aliquotation, storage conditions and duration | Y | Supplementary Information: Homogeneity and stability assessments: p3-5 (SM1) and p9-11 (SM2) |
| **2. NUCLEIC ACID EXTRACTION** | | |
| Description of extraction method including amount of sample processed | N | Not applicable (N/A): no extraction was performed |
| Volume of solvent used to elute/resuspend extract | N | N/A (no extraction was performed) |
| Number of extraction replicates | N | N/A (no extraction was performed) |
| Extraction blanks included? | N | N/A (no extraction was performed) |
| **3. NUCLEIC ACID ASSESSMENT AND STORAGE** | | |
| Method to evaluate quality of nucleic acids | Y | Supplementary Information Figure S2 (SM1); Figures S3-4 (SM2). |
| Method to evaluate quantity of nucleic acids (including molecular weight and calculations when using mass) | Y | Reference value assignment: Supplementary Information Table S1 (SM1); Table S10 (SM2) |
| Storage conditions: temperature, concentration, duration, buffer, aliquots | Y | Supplementary Information page 3 (SM1), page 8 (SM2) |
| Clear description of dilution steps used to prepare working DNA solution | Y | Each participating laboratory determined the dilution of the study materials and volume added per dPCR. Information available upon request. |
| **4. NUCLEIC ACID MODIFICATION** | | |
| Template modification (digestion, sonication, pre-amplification, bisulphite etc.) | Y | Supplementary Information. Plasmid restriction digestion: page 2 (SM1), page 8 (SM2); wild-type DNA sonication (SM1) page 3. |
| Details of repurification following modification if performed | N | N/A |
| **5. REVERSE TRANSCRIPTION** | | |
| cDNA priming method and concentration | N | N/A |
| One or two step protocol (include reaction details for two step) | N | N/A |
| Amount of RNA added per reaction | N | N/A |
| Detailed reaction components and conditions | N | N/A |
| Estimated copies measured with and without addition of RT* | N | N/A |
| Manufacturer of reagents used with catalogue and lot numbers | N | N/A |
| Storage of cDNA: temperature, concentration, duration, buffer and aliquots | N | N/A |
| **6. dPCR OLIGONUCLEOTIDES DESIGN AND TARGET INFORMATION** | | |
| Sequence accession number or official gene symbol | Y | Materials and Methods: Study Materials section/page 6 |
| Method (software) used for design and *in silico* verification | Y | Participating laboratories designed or applied published methods as per Supplementary Information Tables S5-S6. Further information available upon |
| Location of amplicon | Y | Figure 2 |
| Amplicon length | Y | Supplementary Information Tables S11-S12 |
| Primer and probe sequences (or amplicon context sequence)** | Y | Supplementary Information Tables S11-S12 |
| Location and identity of any modifications | Y | Supplementary Information Tables S11-S12 |
| Manufacturer of oligonucleotides | Y | Supplementary Information Tables S11-S12 |

| 7. dPCR PROTOCOL | | |
|---|---|---|
| Manufacturer of dPCR instrument and instrument model | **Y** | Supplementary Information Tables S13-S14 |
| Buffer/kit manufacturer with catalogue and lot number | **Y** | Supplementary Information Tables S13-S14 (lot number available upon request) |
| Primer and probe concentration | **Y** | Supplementary Information Tables S11-S12 |
| Pre-reaction volume and composition (incl. amount of template and if restriction enzyme added) | **Y** | Supplementary Information Tables S13-S14 (amount of template information available upon request) |
| Template treatment (initial heating or chemical denaturation) | **N** | Not performed |
| Polymerase identity and concentration, Mg++ and dNTP concentrations*** | **N** | Manufacturer proprietary information |
| Complete thermocycling parameters | **Y** | Supplementary Information Tables S11-S12 |
| **8. ASSAY VALIDATION** | | |
| Details of optimisation performed | **N** | Available upon request |
| Analytical specificity (vs. related sequences) and limit of blank (LOB) | **N** | Available upon request |
| Analytical sensitivity/LoD and how this was evaluated | **N** | Available upon request |
| Testing for inhibitors (from biological matrix/extraction) | **N** | Available upon request |
| **9. DATA ANALYSIS** | | |
| Description of dPCR experimental design | **N** | Available upon request |
| Comprehensive details negative and positive of controls (whether applied for QC or for estimation of error) | **N** | Available upon request |
| Partition classification method (thresholding) | **N** | Available upon request |
| Examples of positive and negative experimental results (including fluorescence plots in supplemental material) | **N** | Available upon request |
| Description of technical replication | **N** | Available upon request |
| Repeatability (intra-experiment variation) | **Y** | Each participating laboratory reported measurement uncertainties which included assay repeatability (Supplementary Information S15-S16) |
| Reproducibility (inter-experiment/user/lab etc. variation ) | **Y** | Table 1 and 2 |
| Number of partitions measured (average and standard deviation ) | **Y** | Supplementary Information Tables S13-S14 |
| Partition volume | **Y** | Supplementary Information Tables S13-S14 |
| Copies per partition (λ or equivalent ) (average and standard deviation) | **N** | Available upon request |
| dPCR analysis program (source, version) | **Y** | Supplementary Information Tables S13-S14 |
| Description of normalisation method | **N** | N/A |
| Statistical methods used for analysis | **N** | Available upon request |
| Data transparency | raw data available on request: | |

**Table S1.** dMIQE2020 checklist for authors, reviewers and editors. Authors should fill detail whether information is provided. Where 'yes' is selected use comment box to detail location of information or to include the information. Where 'no' is selected use comment box to outline rationale for omission. Sections 4 and 5 may not apply depending on experiment.

* Assessing the absence of DNA using a no RT assay (or where RT has been inactivated) is essential when first extracting RNA. Once the sample has been validated as DNA-free, inclusion of a no-RT control is desirable, but no longer essential.

** Disclosure of the primer and probe sequence is highly desirable and strongly encouraged. However, since not all commercial pre-designed assay vendors provide this information when it is not available assay context sequences must be submitted (Bustin et al. Primer sequence disclosure: A clarification of the miqe guidelines. Clin Chem 2011;57:919-21.)

*** Details of reaction components is highly desirable, however not always possible for commercial disclosure reasons. Inclusion of catalogue number is essential where component reagent details are not available.
